# inPOSE: a flexible toolbox for chromosomal cloning and amplification of bacterial transgenes

**DOI:** 10.1101/2021.10.28.466346

**Authors:** Ranti Dev Shukla, Ágnes Zvara, Ákos Avramucz, Alona Yu. Biketova, Akos Nyerges, László G. Puskás, Tamás Fehér

## Abstract

Cloning genes and operons encoding heterologous functions in bacterial hosts is almost exclusively carried out today using plasmid vectors. This has multiple drawbacks, including the need for constant selection and the variation in copy numbers. Chromosomal integration of transgenes has always offered a viable alternative, however, to date it has been of limited use due to its tedious nature and to being limited often to a single copy. We introduce here a strategy that uses bacterial insertion sequences, the simplest autonomous transposable elements to insert and amplify genetic cargo into a bacterial chromosome. Transgene insertion can take place either as transposition or homologous recombination, and copy-number amplification is achieved using controlled copy-paste transposition. We display successful use of IS*1* and IS*3* for this purpose in *Escherichia coli* cells, using various selection markers. We demonstrate the insertion of selectable genes, an unselectable gene, and a five-gene operon in up to two copies in a single step. We continue with the amplification of the inserted cassette to double-digit copy numbers within two rounds of transposase induction and selection. Finally, we analyze the stability of the cloned genetic constructs in the lack of selection and find it to be superior to all investigated plasmid-based systems. Due to the ubiquitous nature of transposable elements, we believe that with proper design, this strategy can be adapted to numerous further bacterial species.

## Introduction

Many biologists today link the birth of recombinant DNA technology to the first successful insertion of a gene into a plasmid vector using restriction digestion and ligation [1]. This process, now referred to as gene cloning has not only maintained its importance in the field, but has also gone through numerous technical improvements to increase its efficiency, speed and flexibility [2]. Today, the fastest and simplest way to maintain and/or express a gene or operon of interest is to clone it into a bacterial plasmid, usually choosing one capable of replication in *Escherichia coli*. Protein expression from plasmids, however, may significantly alter the physiology of the host cell, especially in the case of high copy-numbers, thereby derailing fundamental studies [3, 4]. Plasmids also require constant selection, often warranted by antibiotics and their respective resistance genes. From an industrial viewpoint, this not only poses an additional cost due to the administration of the antibiotic, but also due to the need of its removal from the final fermentation product. In addition, secreted resistance factors, like the β-lactamase enzyme allow plasmids to be lost from cells during selection, leading to inhomogeneous cell populations [5]. A similar impairment of experimental reproducibility can be caused by variations in plasmid copy numbers [6–8]. These factors all indicate the necessity to develop techniques permitting transgene insertion into the bacterial chromosome.

The intent to modify bacterial genomes is probably as old as our knowledge concerning DNA and genes, and as soon as the first DNA sequences became available, precisely targeted genome editing became a realistic goal. The initial technique widely used for this purpose relied on conditionally replicative plasmids (a.k.a. suicide plasmids) and the RecA recombinase enzyme of the host cell [9–12]. In a two-step process of homologous recombination, the allele on the genome was exchanged to the allele carried by the suicide plasmid. Later, linear DNA fragments flanked by appropriate homology boxes turned out to be similarly applicable for genome editing [13, 14]. The invention of the polymerase chain reaction (PCR)[15] further aided this technique by simplifying linear DNA construction [16]. Such fragments, when electroporated into target cells, could act as the donor DNA in single-step gene-exchange reactions [17]. The key enzymes facilitating this reaction were inducible phage-derived recombinases: the λ-Red system was probably the most popular set used, but numerous other viral recombinases have become available for this purpose since then [18–20]. This technique, often referred to as recombineering, quickly became dominant due to its efficiency in generating point mutations, large deletions and small insertions [21–23]. Making large (> 3 kbp) insertions was also possible if the targeted genomic site was cleaved during the recombination process [24]. This however, required the insertion of a landing pad first, which contained the target site of the homing endonuclease responsible for the *in vivo* chromosomal cleavage. The discovery of the CRISPR-Cas system, and its application as a programmable RNA-directed endonuclease [25, 26] made cleavage-assisted recombination generally applicable without the prior integration of a landing pad [27].

Insertion sequences (IS elements or ISes) are the smallest autonomous mobile genetic elements found in nature [28]. They comprise only one or two genes, responsible for the transposition process, surrounded by inverted repeats. To date, 29 families of ISes have been described [28]. ISes belong to prokaryotic transposable elements (TEs), along with more complex members like transposons, integrative conjugative elements, integrative mobilizable elements, type I and II introns, transposable and satellite prophages, mobile genomic islands, inteins, retrons and IStrons [29, 30]. Bacterial TEs have made a significant contribution to reshaping the genomes of their hosts over evolutionary time scales [31, 32], and the effect of their mobility is readily observed in everyday medical microbiology. Examples of the latter include altering the virulence of pathogens [33], mobilizing antibiotic resistance genes [34, 35] or inducing antibiotic resistance [36].

Bacterial transposons differ from ISes in the sense that in addition to transposases, they also contain genes encoding functions unrelated to transposition. Over the decades, transposons have become popular tools of molecular biology, used for gene delivery, mutagenesis and functional genomics studies [37]. Perhaps the most generally known is Tn*5*, which has been applied to knock-in genes, create mutant libraries, study gene essentiality or create reduced genomes [38, 39]. Examples utilizing further transposons are nevertheless countless: random transposon insertions of Mu, Tn*3*, Tn*5* and Tn*1000* have been used for priming sequencing libraries [40]. Members of the Tn*3* family have been employed in *Escherichia coli* for in-frame tagging of gene libraries [41] or pentapeptide-scanning mutagenesis of single genes cloned in plasmids [42]. Due to its high target-selectivity, wild type Tn*7* is appropriate for gene transfer in-between replicons [43], but modified Tn*7* lacking site-specificity is also available for random mutagenesis studies [44]. Lately, two natural examples of Tn*7*-related transposons were identified in *Vibrio cholerae* [45] and *Scytonema hofmanni* [46], respectively, both of which carry various elements of the CRISPR-Cas machinery. These TEs allow RNA-directed insertion of the DNA-transposons, warranting the possibility of programmable gene delivery at predefined sites of bacterial genomes using transposition. More recently, in vitro application of the Tn*5* transposase together with next generation sequencing technology has opened multiple avenues in genomics studies, including the analysis of 3D genomic structure, detection of copy number variations, sequencing long fragments and mapping DNA-methylation (reviewed by [47]).

As opposed to transposons, ISes are much less often used as molecular biology tools. Notable cases nevertheless exist: for example, IS*21* has been used for linker-scanning mutagenesis of an enzyme-encoding gene cloned in *E. coli* [48]. Another exciting work described the reprogramming of IS*608* (IS*Hp608*) of *Helicobacter pylori* to allow predictable integration at chosen target sites, both in vitro and in vivo, years before the outbreak of the CRISPR-era [49]. Similarly, fusion of the IS*30* transposase to various specific DNA-binding proteins permitted the directed integration of the IS*30* element both in *Salmonella* Enteritidis [50] and in zebrafish [51]. A method relying on IS*608* insertion by homologous recombination, followed by its precise transposase-mediated excision has also been developed for editing short genomic segments [52].

In this work, we test the application of two IS elements, IS*1* and IS*3* for chromosomal gene cloning and amplification in *E. coli*. We report the insertion of marked and unmarked genes into genomic ISes in one or two copies, followed by the copy number amplification of the loaded ISes exploiting their copy-and-paste transposition.

## Methods and Materials

### Strains, chemicals and media

The *E. coli* strains modified in this study are listed in Table 1. Bacteria were grown in liquid Luria Bertani medium (LB), or on LB plates containing 1.5% agar. Components of the media were obtained from Molar Chemicals Kft., Halásztelek, Hungary. Antibiotics were obtained from Sigma Aldrich (St. Louis, MO, USA) and were used in the following concentrations: chloramphenicol (Cm): 25 μg/ml, ampicillin (Ap): 100 μg/ml, kanamycin (Km): 25 μg/ml, spectinomycin (Sp): 50 μg/ml, anhydrotetracycline (aTc): 50 ng/ml. Antibiotic-gradient plates were made applying the protocol of Szybalski and Bryson [53], but using 60x higher Sp and 20x higher Km concentrations in the top layer, compared to the values listed above. Plasmid preparations were made using the Zippy Plasmid Mini Prep Kit (Zymo Research Ltd., Orange County, CA, USA). Horizontal electrophoresis of DNA was carried out using 1% Seakem LE agarose gels (Lonza, Basel, Switzerland). All cloning and molecular biology experiments were carried out according to established protocols, unless otherwise noted [54].

**Table 1.** **E. coli** strains modified in this study

| *E. coli* strain | Reference | GenBank Acc. No. |
| --- | --- | --- |
| MG1655 | [55] | NC_000913.3 |
| DH5a | [56] | NZ_CP080399 |
| MDS16 | this work | - |
| MDS27 | this work | - |
| MDS30 | this work | - |
| MDS39R2 | this work | - |
| MDS42 | [57] | AP012306 |
| BL21(DE3)pLysE | [58] | NC_012947.1 |
| BLK09 | [59] | CP014641 |
| BLK16 | [59] | CP014642 |

### Plasmids used in this study

Plasmids pSG76-CS (GenBank: AF402780.1) [23] pST76-A (GenBank: Y09895.1), pST76-K (GenBank: Y09897.1), pSG76-K (GenBank: Y09894.1) [60], pZA31CFPtetR [61], pORTMAGE2 and pORTMAGE4 have been described previously [62]. Plasmid pSG78-A is identical to pSG76-A (GenBank: Y09892.1), except for containing an additional P-SceI site next to the I-SceI site. Plasmid pCDM4 was a kind gift of Prof. Mattheos Koffas [63]. Plasmid pUTLIQ_vioABCDE was a kind gift of Prof. Kanji Nakamura. Plasmids pKDsg-ack and pCas9cr4 were kind gifts of Prof. Kristala Prather [27].

To make plasmid pKDsg-IS1, the pKDsg-ack plasmid was amplified with primers pKD1 and IS1noScarFwd as well as pKD2 + IS1noScarRev using Q5 DNA polymerase (New England Biolabs, Ipswitch, MA, USA). Both products were DpnI digested and cleaned with the Viogene Gel/PCR DNA Isolation Kit (Viogene-Biotek Corporation, New Taipei, Taiwan R.O.C.). These fragments were assembled in a Circular Polymerase Extension Cloning (CPEC) [64] reaction using Phusion DNA polymerase (New England Biolabs, Ipswitch, MA, USA) and transformed into *E. coli* MDS42 fast competent cells and selected on Sp plate at 30 °C. Colony PCR-screening was done with primers pKDSeq5 + IS1noScarCheckR. Plasmid was purified from a PCR-positive colony and the pKdSeq5+pKDseq3 PCR fragment amplified from the plasmid was sequenced with primer pKDseq5 for verification. Plasmid pKDsg-IS3 was made similarly, except that primers IS1noScarFwd, IS1noScarRev and IS1noScarCheckR were replaced by IS3noScarFwd, IS3noScarRev and IS3noScarCheckR, respectively.

To generate pST76AIS1X, two overlapping PCR fragment were amplified from the IS*1* element of the *E. coli* MDS42_IS1 genome with IS1_Ssp_up + IS1_Xho_Rev and IS1_Xho_Fwd + IS1_Ssp_Dn, respectively. Both fragments were cleaned with the Viogene Gel/PCR DNA Isolation Kit (Viogene-Biotek Corporation, New Taipei, Taiwan R.O.C.) and then fused in an overlapping PCR to make an IS*1* fragment lacking inverted repeats but having SspI sites on the ends and an XhoI site in the center. The PCR product was treated with T4 PNK enzyme (Thermo Fisher Scientific, Waltham, MA, USA) for phosphorylation and was gel extracted with the Viogene Gel/PCR DNA Isolation Kit. The pST76-A plasmid was cut with fast digest BsaI (New England Biolabs, Ipswitch, MA, USA) followed by phosphatase treatment (Fast AP, Thermo Fisher Scientific). Linear pST76-A plasmid was ligated with the PCR fragment using T4 ligase enzyme overnight at 16 °C followed by heat inactivation at 65 °C for 10 minutes and dialysis for 1 h on a 50 nm pore-sized membrane (Millipore). The ligation product was transformed into *E. coli* MDS42 fast competent cells and colonies were selected overnight at 30 °C on Ap plates. Colony PCR was done using primers pSTA-F + pSTA-R to find positive clones. Two positive clones were used to make plasmid preparations, and the pSTA-F + pSTA-R fragment was PCR-amplified for their sequencing using the T7 universal primer. Plasmid pST76AIS3X was made similarly, except that the IS*3* element of the *E. coli* MDS30 genome was the amplified as two overlapping PCR fragments using primers IS3_Ssp_up + IS3_Xho_Rev and IS3_Xho_Fwd + IS3_Ssp_Dn.

To obtain plasmid pST76AIS1X::VioABCDE, the pST76AIS1X plasmid was amplified using primers IS1VioFwd and IS1VioRev. The violacein operon was amplified as two overlapping PCR fragments from plasmid pUTLIQ_vioABCDE using primers pUTLIQfwd + VioCendRev2 and VioCendfw + pUTLIQRev7, respectively. All the three fragments were assembled uisng the NEBuilder^®^ HiFi DNA Assembly Cloning Kit (New England Biolabs, Ipswitch, MA, USA) and transformed into *E. coli* MDS42 fast competent cells. Colony PCR-screening was done with primers pSTA-F + VioARev. Plasmids were prepared from positive clones and were verified by restriction digestion using the SspI fast digest enzyme (Thermo Fisher Scientific, Waltham, MA, USA). Plasmid pST76AIS3X::VioABCDE was made similarly, except that the the initial PCR was amplified from plasmid pST76AIS3X using primers IS3VioFwd and IS3VioRev.

Plasmid pST76AIS1X::VioABCDE_SpR was generated by inserting an SpR gene into pST76AIS1X::VioABCDE by recombineering. This required the initial amplification of the SpR gene from the PCDM4 plasmid using primers (SpRVioIntFw and SpRVioIntRev) that carried the appropriate homology boxes to direct the integration of the PCR fragment into the region between the VioE gene and 3’ fragment of IS*1*. This PCR fragment was transformed into *E. coli* MDS42/pORTMAGE4/pST76AIS1X::VioABCDE induced by a transient heat shock (see below) in order to express the λ-Red recombinases to facilitate homologous recombination. Cells were plated on Ap+Sp agar plates at 30 °C. Colony PCR-screening was done with primer pairs pSTA-R + SmRev as well as TetR1 + SmRev, respectively. Double positive colonies were replica-plated on Sp as well as Cm plates in parallel. Sp^R^Cm^S^ colonies were picked for plasmid preparation. Plasmids were verified with double digestion using KpnI + NheI fast digest enzymes (Thermo Fisher Scientific, Waltham, MA, USA).

Plasmid pST76AIS1X::VioABCDE_KmR was made in a way similar to pST76AIS1X::VioABCDE_SpR except that the KmR gene to be integrated was amplified from plasmid pSG76-K using primers KmRVioIntFw and KmRVioIntRev. Colony screening was done with primer pairs pSTKF + pSTAR, pSTKF + TetR1 and Km_HindIII_Fwd + Km_HindIII_Rev. Triple positive clones were replica-plated on Km as well as Cm plates in parallel. Km^R^Cm^S^ colonies were picked for plasmid preparation. Plasmids were digested using PvuII fast digest enzyme (Thermo Fisher Scientific, Waltham, MA, USA) and compared with the digestion pattern of the pST76AIS1X::VioABCDE plasmid.

The pST76AIS3X::VioABCDE_SpR plasmid was made like pST76AIS1X::VioABCDE_SpR, except that the recombineering took place in *E. coli* MDS42/pORTMAGE4/pST76AIS3X::VioABCDE.

Plasmid pST76AIS3X::VioABCDE_KmR was generated similarly to pST76AIS1X::VioABCDE_KmR except that the recombineering took place in *E. coli* MDS42/pORTMAGE4/pST76AIS3X::VioABCDE. In the end, the PvuII-digested plasmid preparations were compared to PvuII-digested pST76AIS3X::VioABCDE plasmid.

To make pZA31insAB’tetR plasmid, the *insA* ORF of IS*1* was PCR-amplified from the *E. coli* MG1655 genome with Q5 DNA polymerase using the InsAfw and InsArev primers. The *insB* of IS*1* was PCR-amplified similarly using the InsBfw + InsBrev primers. The two products were fused in an overlap-PCR to make InsAB’ linear fragment, carrying the the two ORFs of IS*1* within the same frame. This fragment was digested with SphI + BamHI Fast Digest enzymes (Thermo Fisher Scientific, Waltham, MA, USA), and the product was purified with the Viogene Gel/PCR DNA Isolation Kit. The pZA31CFPtetR vector was PCR-amplified with primers pZA31rev + pZA31fw, digested with SphI + BamHI and ligated with the digested InsAB’ fragment using the T4 ligase enzyme. The ligation product was transformed into *E. coli* MDS42 followed by selection on Cm agar plate. The obtained colonies were PCR-screened with primers tetP5 + IS1/1. Clones displaying a PCR product of 700 bp were used to make plasmid preparations which were sequenced using the tetP5 primer.

To make pSTinsAB’tetR plasmid (carrying a temperature sensitive origin of replication), the pSC101* origin + *repA* gene were PCR amplified together from plasmid pST76-K using primers pSToriF + pSToriR followed by DpnI digestion and purification. Both the PCR product and plasmid pZA31insABtetR were digested with XmaJI + AseI (Thermo), purified, and ligated using the T4 ligase. The ligation product was transformed into *E. coli* MDS42 followed by selection on Cm agar plate at 30 °C. Colonies were PCR-screened with primers pSTA-R + tetP5. The temperature sensitive nature of PCR-positive colonies was verified by the absence of their growth at 37 °C.

To generate plasmid pZA31tnp3tetR, the codon-optimized ORFs of IS*3* were ordered to be synthesized by Eurofins Genomics GmBH. These two ORFs, *insE* and *insF* were synthesized to be in the same reading frame, and the entire construct was flanked by SphI and BamHI sites. The synthetic *insEF*’ gene was cloned into the SphI+BamHI sites of pZA31CFPtetR, as described above. Colony-PCR screening was carried out with the primer pairs tetP5 + tnp3rev and tetP5 + pZArev, respectively.

To make pSTtnp3tetR plasmid (carrying a temperature sensitive origin of replication), the pSC101* origin was inserted exactly as described in the case of pSTinsAB’tetR above.

To make pSG78A_full_IS3::SpR plasmid, the IS*3*::SpR cassette was PCR-amplified from the *E. coli* MDS30_IS*3*::SpR genome (generated by recombineering, see below) with primers Ecob1028fw + Bamb1028rev. The PCR fragment was DpnI digested, purified and sequenced from both ends. After confirming the correct sequence, the PCR fragment and the pSG78-A vector were both digested with EcoRI + BamHI, purified and ligated together using T4 ligase. The ligation product was heat-inactivated at 65 °C, dialyzed on a 50 nm pore-sized membrane (Millipore) and transformed into MDS42-π fast competent cells (expressing the Pir protein) followed by overnight selection on Ap + Sp plate at 37°C. Colonies were PCR-screened with pSGR + SmFwd primers. Two positive clones were used for plasmid preparation followed by plasmid re-transfromation into MDS42-π to ensure plasmid purity. Three colonies from each plate were picked, cultured in 5 ml of LB + Ap + Sp, ovrnight at 37 °C, then the culture was PCR-amplified with three primer pairs (Ap1+ SmRev, pSGR + SmFwd and Ap1 + pSGR). Two clones displaying correct products were picked for plasmid preparation and one plasmid was sent for sequencing with T7 + pSGR.

Plasmid pCas9IS1, capable of directing Cas9 to cleave IS*1*, was generated from plasmid pCas9 by annealing IS1-SPC1 and IS1-SPC2 single stranded phosphorylated oligonucleotides and ligating into the BsaI site, as described previously in the pCas9 protocol (Addgene #42876)[65]. Colony PCR-screening was done with the pCas9Fwd and IS1_SPC_Chek_Rev primer pair. Plasmid was prepared from positive clones and plasmid PCR was made using primers pCas9Fwd + pCas9Rev for sequence verification of the spacer.

Plasmid pCas9_IS3, capable of directing Cas9 to cleave IS*3*, was generated similarly to pCas9IS1, except that the spacer was made by annealing the following pairs of phosphorylated oligonucleotides: SPC-IS3-Fwd-long and SPC-IS3-Rev-long (for 30 bp-long spacer) or SPC-IS3-Fwd-short and SPC-IS3-Rev-short (for 20 bp-long spacer). Colony PCR-screening was done with the pCas9Fwd and IS3_SPC_Chek_Rev primer pair.

### Rapid electroporation of *E. coli* cells

In our fast protocol to generate electrocompetent cells, *E. coli* cultures were grown in 10 ml of LB medium (harboring the appropriate antibiotic) to an OD600 value of 0.45-0.55. Cells were pelleted at 10,000 g for 2 min, and resuspended in 2×1.5 ml of ice-cold sterile Milli-Q water three times. The final resuspension took place in 40 μl of water. For electroporation, the cell suspension was mixed with the DNA, and the mixture was transferred into electroporation-cuvettes harboring a 1 mm gap (Cell Projects Ltd, Herrietsham, UK). Electroporation was carried out in a MicroPulser electroporator (BioRad, Hercules, CA, USA) set to a voltage of 1.8 kV. The cells were recovered in 1 ml of LB, and shaken for 1-2 h at 30 or 37 °C (depending on the specific case). Finally, 10-100% of the recovery culture was plated on LB Agar plates harboring the appropriate antibiotic.

### Integration of resistance genes into genomic IS elements by recombineering

For linear DNA-mediated recombineering, the PCR fragments targeting IS*1* were made using primers IS1ASpF and IS1ASpR using the spectinomycin-resistant pCDM4 plasmid as a template, or IS1ACmR_F and IS1ACmR_R using the chloramphenicol-resistant pSG76CS as a template. To test the effect of long homologies, primers IS1Sp100F and IS1Sp100R were used to amplify the pCDM4 plasmid. PCR fragments targeting IS*3* were made using primers IS3SpF and IS3SpR using pCDM4 plasmid. For primer sequences, see **Table S1** of the Supplement.

A fully grown overnight culture of the host *E. coli* strain harboring the pORTMAGE2 plasmid was diluted 100-fold into 50 ml LB+Ap. The culture was grown to OD600 of 0.45 to 0.55, then the recombinase genes were induced with a transient heat shock (42 °C, 15 min) followed by a 10 min cooling on ice. This culture was used to make fast electrocompetent cells (see above) which were electroporated with 100 ng of linear PCR fragment (antibiotic resistance gene flanked by homology boxes corresponding to the respective IS element). After 2 h of recovery at 30 °C, 100 μl was spread on either Sp or Km plate (depending on the resistance gene) and incubated overnight at 37 °C. Next day, single co-integrants were screened by colony-PCR using a pair of primers that hybridize to the resistance gene and the chromosomal segment neighboring the targeted IS elements, respectively.

In order to obtain double co-integrants, Cas9 selection was included. The protocol started out as above, but the recovery culture was grown overnight at 30 °C. The next day, it was was added to 50 ml LB and grown to OD600 of 0.45 to 0.55 and then 10 ml fractions of the culture were used to make fast competent cells (see above) and 100 ng of pCas9_IS1 or pCas9_IS3 plasmid was electroporated. After a 2 h recovery in LB at 30 °C, Cm was added to select for the pCas9-derived plasmid and was shaken overnight at 30 °C. On the third day, 100 μl was spread on either Sp+Cm or Km+Cm plates (depending on resistance of the linear cassette) and incubated overnight at 37 °C. On the fourth day, double co-integrants were screened by colony PCR targeting all potential genomic IS elements.

### Integration of a resistance gene or the VioABCDE operon into genomic IS elements using NO-SCAR

A fully grown overnight culture of the host *E. coli* strain harboring plasmids pCas9-Cr4 and either pKDsg-IS1 or pKDsg-IS3 (targeting IS1 and IS3, respectively), was diluted 100-fold into 10 ml LB+Cm+Sp. The culture was grown at 30 °C to OD600 of 0.45 to 0.55 and 0.2% L-arabinose was added (inducing the λ-Red recombinases), followed by further growth at 30 °C for 15 min and cooling on ice for 10 min. This culture was used to prepare electrocompetent cells (see above) which were electroporated with 100 ng of linear PCR fragment (KmR gene flanked by homology boxes corresponding to the respective IS element). After 2 h of recovery at 30 °C, 900 μl was spread on a Km+aTc plate (to induce the CRISPR/Cas system) and incubated overnight at 37 °C. Next day, single co-integrants were screened by colony-PCR using a pair of primers that hybridize to the resistance gene and the chromosomal segment neighboring the targeted IS elements, respectively.

To integrate the VioABCDE operon into genomic IS elements, linear fragments containing the VioABCDE pathway and the resistance marker gene, all flanked by homology boxes corresponding to the targeted IS elements were needed. These were generated by linearizing the pST76AIS3::VioABCDE_KmR plasmid by SspI digestion or linearizing the pST76AIS3::VioABCDE_SpR plasmid by KpnI+MunI+NheI co-digestion. The integration protocol was the same as above, except 0.4% L-arabinose was used for the first induction, 600-800 ng of linear fragment was electroporated, and the recovery culture was spread on a Km+Sp+Cm+aTc plate and incubated overnight at 30 °C.

### Integration of non-selectable genes into genomic IS elements using NO-SCAR

A fully grown overnight culture of the host *E. coli* strain harboring plasmids pCas9-Cr4 and either pKDsg-IS1 or pKDsg-IS3 (targeting IS1 and IS3, respectively), was diluted 100-fold into 10 ml LB+Cm+Sp. The culture was grown at 30 °C to OD600 of 0.45 to 0.55 and 0.4% L-arabinose was added (inducing the λ-Red recombinases), followed by further growth at 30 °C for 15 min and cooling on ice for 10 min. This culture was used to prepare electrocompetent cells (see above) which were electroporated with 100 ng of linear PCR fragment (*gfp* gene flanked by homology boxes corresponding to the respective IS element). After 2 h of recovery at 30 °C, 0.5 μl was spread on a Cm+Sp+aTc plate (to induce the CRISPR/Cas system) and incubated overnight at 37 °C. Next day, single co-integrants were screened by colony-PCR using a pair of primers that hybridize to the resistance gene and the chromosomal segment neighboring the targeted IS elements, respectively.

### Integration of marked IS elements into the genome using transposition

A fully grown overnight culture of the host *E. coli* strain (MDS30 or MDS42) harboring the pSTtnp3tetR plasmid was diluted 100-fold into 5 ml LB+Cm+aTc, and was grown at 30 °C for 2 hrs. Then all the 5 ml of this culture were used to make fast electro competent cells (see above) which were electroporated with 150 to 200 ng of pSG78A_Full_IS3::SpR. After 1 h of recovery at 30 °C, 100 μl was spread on Sp plate and incubated overnight at 37 °C. Next day, several colonies were replica-plated on Cm, Ap and Sp plates at 37 °C. One day later, Sp^R^Cm^S^Ap^S^ colonies (i.e. those that have lost both plasmids but retain the IS*3*::SpR) were inoculated into liquid cultures to generate glycerol stocks. The putative genomic insertions of IS*3*::SpR were localized using ST-PCR (see below).

### Semi-Random Two-Step PCR (ST-PCR)

To localize a known sequence (IS*3*::SpR in our case) within a bacterial genome (*E. coli* MDS30 or MDS42), we applied Semi-Random Two-Step PCR (ST-PCR) [66] with some modifications. Briefly, genomic DNA was PCR-amplified from the colonies of interest first using primer pairs Sp439Rev + CEKG2B and Sp110Fwd + CEKG2B, applying the following program: 94 °C 2 min, followed by 6 cycles of 94 °C 30 sec denaturation, 53 °C 30 sec annealing with 1 °C decrease each cycle, and 72 °C 3 min elongation. The DNA product generated by the first PCR was diluted 5x with TE buffer and subsequently used as a template for nested PCR, with primer pairs SmFwd + CEKG4 and Sp347Fwd + CEKG4 as corresponding to the first two PCRs, respectively. The applied program was 30 cycles of 94 °C 30 sec, 65 °C 30 sec, 72 °C 3 min. The products were separated on 1% agarose gels and bands of interest were extracted (using Viogene Gel/PCR DNA Isolation Kit) and sequenced with SmFwd or Sp347Fwd (depending on which was used for the nested PCR). Relative positions of the primers used for ST-PCR are shown on **Figure S1**.

### Amplification of IS elements marked with antibiotic resistance genes

A fresh overnight culture of the *E. coli* strain harboring the marked IS element was grown at 37 °C. Fast electrocompetent cells were made (as above), transformed with the respective transposase-expressing plasmid (pZA-ST_insAB’ or pZA-STtnp3), spread on Sp+Cm or Km+Cm plate (depending on antibiotic marker in genome) and incubated overnight at 30 °C. A colony was picked and grown as an overnight starter culture in LB using the same antibiotics. The grown culture was diluted 10,000-fold and grown again at 30 °C in LB with antibiotics and the inducer, aTC (an uninduced control was grown in parallel). Dilution and growth were repeated after 24 h. Next the plasmids were cured by growing the culture at 37 °C in LB without antibiotic selection for five days applying a 10,000-fold dilution during each transfer. From the 5^th^ fully grown serial culture, 1 μl was spread on 60x Sp gradient plate or 20x Km gradient plate and incubated at 37 °C for 24 h. The next day, 15 colonies from the high antibiotic-concentration area of the gradient plate were picked and replica-plated on Cm at 30 °C and Sp or Km at 37 °C. After overnight incubation, Cm^S^Sp^R^ or Cm^S^Km^R^ colonies were chosen and grown in liquid LB without selection at 37 °C. The fully grown cultures were saved as glycerol stocks and/or used for violacein quantification or to prepare genomic DNA for ddPCR analysis.

### Quantification of violacein production

To quantify violacein production of liquid *E. coli* cultures, we applied the modified protocol of Zhu et al. [67]. The investigated strain was grown in LB with or without selection (Sp or Km, depending on resistance marker in genome) for 24 h at 37 °C. One ml of the culture was spun at 13,000 rpm for 10 min. After discarding the culture supernatant, 1 ml of DMSO (Molar Chemicals Kft., Halásztelek, Hungary) was added to the pellet. The solution was vortexed vigorously for 30 s to completely solubilize violacein and was centrifuged again at 13 000 rpm for 10 min to remove cell debris. Next, 200 μl aliquots of the violacein-containing supernatant were transferred to a 96-well flat-bottomed microplate (Greiner Bio-One International, Kremsmünster, Austria) making three technical replicates, and the absorbance was recorded at 585 nm using a Synergy2 microplate reader (BioTek, Winooski, VT, USA).

### Monitoring the stability of violacein production

The stability of violacein production was assessed in two different experiments. The first type of experiment monitored the ratio of purple colonies in the lack of antibiotic selection. Overnight starter cultures of the investigated strains were grown in LB with selection (Sp or Km for the genomic co-integrants, Tc for pUTLIQ-carrying strains) at 37 °C. The culture was pelleted and resuspended twice in water to get rid of antibiotics, and plated in appropriate dilutions on solid LB medium to obtain individual colonies. The number of purple colonies and total colonies was counted on each plate, their ratio yielded the “day 0” value of cells expressing violacein. Next, the cultures were diluted 10,000-fold in LB medium and were fully grown without selection at 37 °C. Plating and enumerating the ratio of purple colonies was repeated to yield the ratio of cells expressing violacein for days 1 to 4. Each strain was investigated in three biological replicates.

The second type of experiment monitored violacein production of liquid cultures grown in the lack of selection. Overnight starter cultures of the investigated strains were grown in LB with selection (Sp or Km for the genomic co-integrants, Tc for pUTLIQ-carrying strains) at 37 °C. Next, the culture was diluted 1,000-fold in LB medium and was fully grown without selection at 37 °C. This dilution/growth cycle was repeated three more times. Each day, 1 ml of the fully grown culture was analyzed for violacein content, as described above. Each strain was investigated in three biological replicates.

### Droplet digital PCR

To determine the copy numbers of marked IS elements within the bacterial chromsome, droplet digital PCR (ddPCR) experiments were performed using the EvaGreen protocol of BioRad QX200 Droplet DigitalPCR system (BioRad, Hercules, CA, USA). The template genomic DNA from E.coli strains were purified with the NucleoSpin Microbial DNA kit (Macherey Nagel), and digested with SspI (Thermo Fisher Scientific, Waltham, MA, USA). Final concentrations in reaction mixtures were the following: 1x QX200 EvaGreen Digital PCR Supermix (BioRad, Hercules, CA, USA), 3pg digested E. coli genomic DNA, 200nM combined primer mix in 25ul final volume. Twenty microliter of reaction mixtures was used for droplet generation using QX200 droplet generator. After partitioning, the samples were transferred into a 96-well plates, sealed and put in a T100 Thermal cycler (BioRad, Hercules, CA, USA). The following cycling protocol were used: 95 °C for 10 min, followed by 40 cycles of 94 °C for 30 s and 60 °C for 1 min followed by 5 min at 4 °C, 5 min at 95 °C and finally at 4-degree infinite hold. The droplets were then read in the FAM channels and analyzed using the QX200 reader (Bio-Rad, Hercules, CA, USA). Primers specific for the Km-resistance gene (Km13fw and Km96rev) and the Sp-resistance gene (Sp347fw and Sp439rev) were designed using the ApE software (https://jorgensen.biology.utah.edu/wayned/ape) and ordered from MWG Eurofins Genomics Gmbh (Ebersberg, Germany). Final copy numbers were normalized to the bacterial single copy *lacZ* gene, amplified with primers lacZ110fw and lacZ200rev. All primer sequences are listed in **Table S1** of the Supplement.

### Whole genome sequencing of bacteria

For whole genome shotgun sequencing, genomic library was prepared from four strains (B0, B1, B2 and B3) using the Nextera XT Library Preparation Kit (Illumina, San Diego, CA, USA), according to the manufacturer’s protocol. The libraries were sequenced with Illumina NextSeq 500 sequencers using 2×150 PE sequencing. Reads from all samples were aligned to the *E. coli* BLK09IS3::VioABCDE_SpR reference genome with BWA [68]. To find all insertion sites we used Smith-Waterman algorithm [69] by filtering all reads where the IS3 element’s 3’ 25 nt (TGATCCTACCCACGTAATATGGACA) or 5’ 25 nt (TGTCCACTATTGCTGGGTAAGATCA) sequence or their reverse complements can be found with maximum 2 mismatches, then with Smith-Waterman algorithm selected all those reads which can be aligned only with more than 10 mismatches to the +/− 200 nt region of the original single insertion site. Again, using the Smith-Waterman algorithm we trimmed all nt from each read which can be aligned to the insertion element. The resulting fastq file contained all reads marking the neighboring sequences of each insertion. This fastq file was remapped by BWA to the *E. coli* BLK09IS3::VioABCDE_SpR reference genome. To mark independent insertion sites, we searched for peak positions different from those of sample B0. To calculate the overall number of independent insertions and repeated re-insertions, relative coverage was measured in the complete alignments by calculating the ratio of reads at +/−10 nts from peak borders.

## Results

The general strategy tested in this work is to use ISes as landing pads to integrate transgenes into bacterial genomes, followed by their copy-number amplification using transposition. There are several considerations supporting this idea: i) to the best of our knowledge, ISes have never been found to be essential, ii) ISes are near-ubiquitous, having been detected in 76% of bacterial genomes [70], iii) the type, number and exact position of ISes in each sequenced genome are well known, iv) copy-and-paste type of ISes have been identified and v) the mobility (transposition rate) of various IS types is often known, allowing the anticipation of potential transgene amplification. We chose to test two copy-and-paste type of IS elements, IS*1* and IS*3*, which are at the high-end and low-end of the transpositional activity scale in *E. coli*, respectively [71]. Concerning the copy numbers of these elements, wild type (wt) *E. coli* K-12 MG1655 (GenBank Acc. No. NC_000913.3) carries 8 copies of IS*1* and 5 copies of IS*3*, while *E. coli* BL21(DE3)pLysGold (NC_012947.1) carries 28 and 4 copies, respectively. In the course of an earlier project carried out in our laboratory however, we deleted all ISes from MG1655 [57], and saved intermediate strains carrying 2 or 1 copies of IS*1* (called MDS39R2 and MDS42IS1, respectively), or 3, 2 or 1 copies of IS*3* (called MDS16, MDS27 and MDS30, respectively). Similarly, during the course of BL21(DE3) genome reduction [59], we generated BLK09 which carries 2 active copies of IS*3*, and BLK16, where the same 2 copies of IS*3* are inactivated by nonsense mutations within the transposase gene. We were therefore in the fortunate situation that we could choose multi-deletion strains carrying limited copies of the targeted elements, thereby simplifying the analysis of transgene integration.

### Targeting ISes by recombineering

In the first type of experiment, we asked whether targeting IS*1* and IS*3* were just as straightforward as targeting other nonessential genes of *E. coli*. We generated linear dsDNA fragments by PCR that harbored an antibiotic resistance gene flanked by two appropriate homology boxes (provided by the PCR primers). We expressed the λ-Red recombinases from the inducible pORTMAGE2 plasmid to facilitate the double-crossover between the respective chromosomal IS element and the resistance-cassette. In parallel experiments, we targeted the single IS*1* copy of MDS42IS1 and the single IS*3* copy of MDS30. As expected, the colonies obtained on selective plates were genomic co-integrants: using colony PCR, 10 out of 10 colonies tested positive in both the IS*1*-targeting and the IS*3*-targeting experiment. Therefore, integrating selective markers into IS elements by recombineering is not different than targeting other genes. It is noteworthy however, that using SpR as a selection marker permitted more than 20-fold higher absolute recombination efficiencies than using CmR (15.26 colony/ng DNA vs. 0.718 colony/ng DNA, respectively, as listed in **Table S2**).

In the second class of experiments, we repeated the recombineering process as above, but used strains carrying two copies of the targeted ISes: MDS39R2 for IS*1* and BLK09 or BLK16 for IS*3*. All colonies obtained on the selective plates were PCR-screened for integration at both loci. On the one hand, when targeting IS*1* in MDS39R2, only 13 of 25 colonies (52%) carried the SpR in either of the IS*1* elements, for the remaining 12, the SpR cassette integrated at an unknown locus. The majority of the known integrations occurred in the YeaJ::IS*1*, the minority in the ais::IS*1* (11 vs 2, respectively) (See **Figure S2** for an example). On the other hand, when targeting IS*3* in BLK09 and BLK16, the resistance cassette was always detectable at either one of the two loci. Again, there was a bias favoring the integration into the IS*3* at locus 1 (gene ECBD2567) vs. locus 2 (gene ECBD0875) (7:3, respectively). The ratios of integration at each locus are shown on **Figure 1**. Double co-integrants were never obtained this way in either the IS*1* or the IS*3*-targeting experiments. When targeting the two copies of IS*1*, we confirmed the observation that using SpR as a transgene yields more co-integrants than using CmR (16.73 vs. 0.33 colony/ng, respectively, see **Table S2**), explaining our preference for SmR in further experiments.

**Figure 1.**
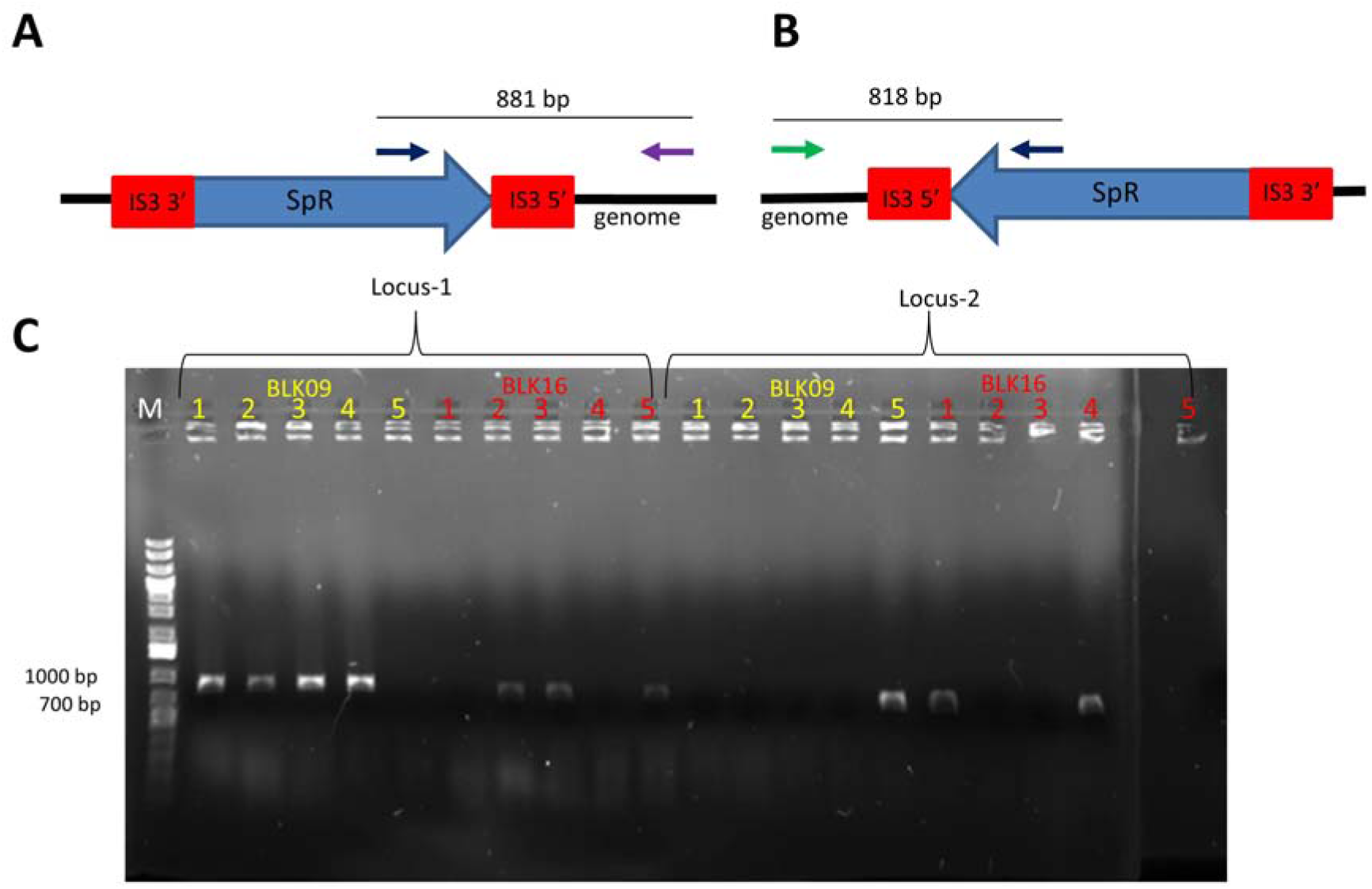
Colony PCR to verify chromosomal SpR integration after recombineering. (A) Expected genetic arrangement and PCR product in case of successful SpR integration at locus-1. (B) Expected genetic arrangement and PCR product in case of successful SpR integration at locus-2. Black arrow: primer SmFw; purple arrow: primer IS3flanking1; green arrow: primer IS3flanking2. SpR: spectinomycin resistance gene. Red boxes depict IS*3*-homologies. (C) Gel electrophoresis of colony-PCR reactions made for screening of 5 *E. coli* BLK09 and 5 *E. coli* BLK16 colonies obtained upon recombineering. Primer pairs either specific for locus 1-integration or for locus 2-integration were used in both cases. M: GeneRuler 1 kb Plus DNA Ladder (Thermo Scientific).

In our third group of experiments, we attempted the integration of the 7.3 kbp-long VioABCDE cassette into the bacterial genome using λ-Red-mediated recombineering. The VioABCDE operon originates from *Chromobacterium violaceum*, and is responsible for producing the purple pigment violacein [72]. Since generating linear VioABCDE fragments (flanked by appropriate homologies) using PCR was unsuccessful, we cloned the operon into the temperature-sensitive pST76-A vector, flanked by homology arms corresponding to IS*1* or IS*3* elements, making pST76AIS1X::VioABCDE and pST76AIS3X::VioABCDE, respectively. (Note that neither of these plasmids contains the inverted repeats of the targeted IS.) Linear DNA fragments generated from these plasmids were substrates of recombineering using pORTMAGE2. These cassettes lacked an antibiotic selection marker, our rationale was to obtain a lawn of cells on nonselective plates, and visually screen for purple sectors. Such purple sectors were never visible by the naked eye, indicating that the rate of recombination is too low for a visual screening strategy. We therefore went on to include an antibiotic resistance gene (SpR or KmR) inside the cassette next to the Vio operon, within the segment flanked by IS-specific homologies. Four such plasmids were generated: pST76AIS1X::VioABCDE_SpR, pST76AIS1X::VioABCDE_KmR, pST76AIS3X::VioABCDE_SpR and pST76AIS3X::VioABCDE_KmR. Again, linear DNA cassettes for recombineering were generated from these plasmids by restriction digestion. When electroporating these cassettes, we obtained mixed results: on the one hand, if targeting genomic IS*1* in MDS42IS1, correct recombination events could be verified, but none of the co-integrant colonies were purple. This was the case despite the fact that all four plasmids used to generate the cassettes granted their host a dark purple colony phenotype. On the other hand, when targeting the IS*3* of MDS30, the PCR-verified genomic co-integrants displayed a purple color: Three out of nine colonies (33%) were PCR-positive when using SpR, and 9/10 (90%) were positive when using KmR as a selection marker (**Figure S3**).

Targeting *E. coli* strains having two copies of IS*3* within the genome also provided the integration of the VioABCDE_SpR cassette with rates acceptable for practical purposes: the fractions of colonies harboring single co-integrants in *E. coli* BLK09, BLK16 and MDS27 strains fall between 20 and 75% (**Table 2**).

**Table 2.** The fraction of colonies obtained on the selective plate that carried the co-integrant at either IS*3* locus, as verified by PCR

|  | Selection marker used |  |
| --- | --- | --- |
|  | SpR | KmR |
| BLK09 | 20% | 65% |
| BLK16 | 75% | 50% |
| MDS27 | 25% | 45% |
| MDS16 | ND | 55% |

Double co-integrants were never obtained this way. We have nevertheless demonstrated that the 9074 bp and 9359 bp-long VioABCDE-SpR and VioABCDE-KmR cassettes, respectively, could be targeted into genomic IS*3* elements by recombineering. In both experiments targeting IS*3* of BLK09 or BLK16, the majority (>90%) of the colonies displayed a purple color, indicating a relatively low rate of false positive resistance. In strain MDS27 however, <50% of colonies were purple to the naked eye, and some of the white colonies were true co-integrants verified by PCR. This exemplifies the strain-dependence of the ratio of colonies with functional expression of the inserted operon. The bias of the integration seen for resistance genes was also present when inserting the five-gene operon, although to a variable extent (**Figure 2**).

**Figure 2.**
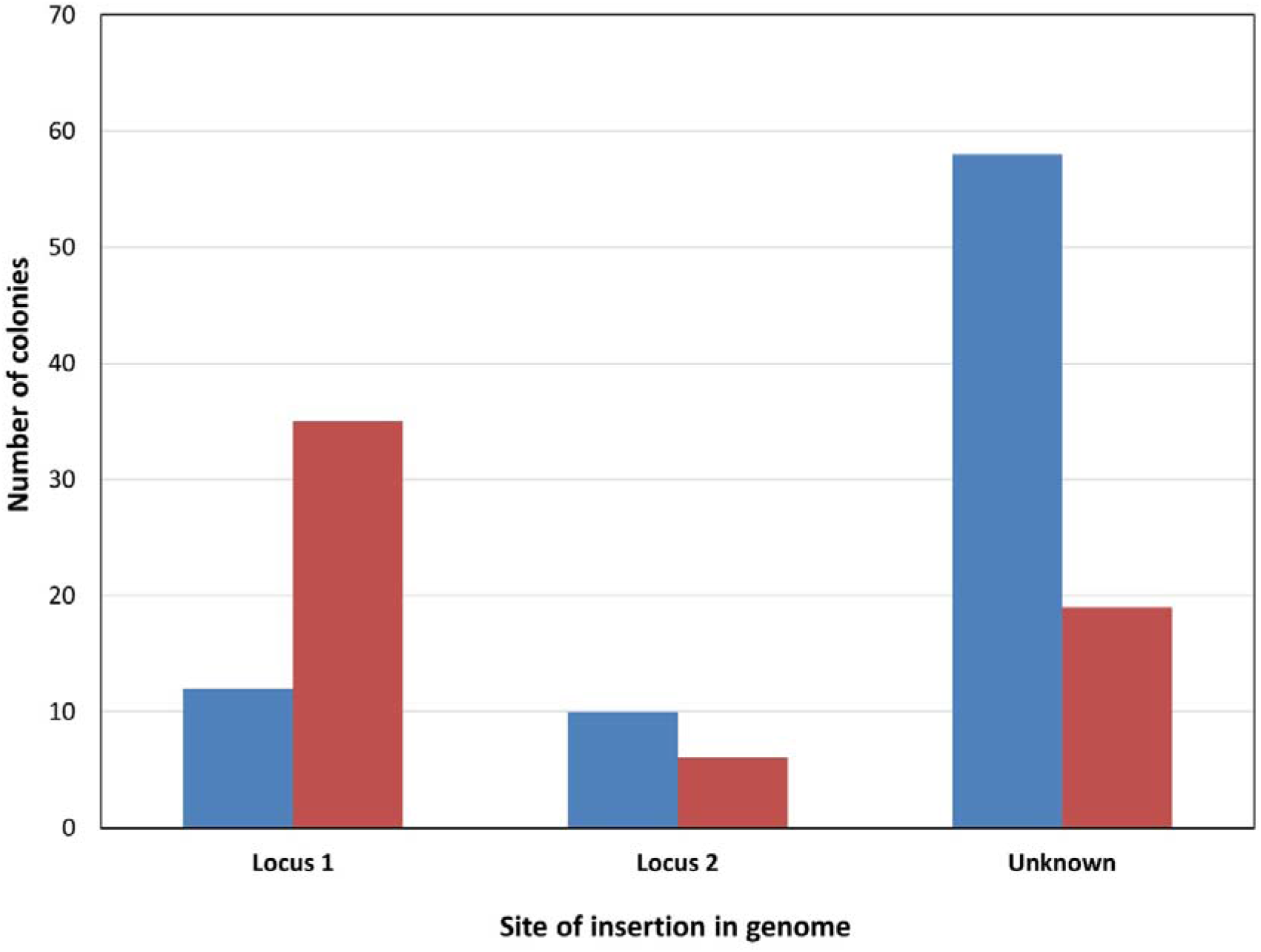
Combined result of recombineering experiments integrating the VioABCDE_Km or VioABCDE_SpR cassette into a chromosomal IS*3*. The number of colonies harboring a PCR-verified integration at locus 1, locus 2, or displaying no PCR-verified integration (“unknown”) are shown for *E. coli* BLK09 (blue) and *E. coli* BLK16 (red).

When targeting *E. coli* MDS16, harboring three IS*3* elements, we managed to detect single co-integrants at all three loci, although we only found one insertion at b0374::IS*3* out of 20 colonies tested (**Figure 3**). We have therefore shown that a relatively long (>9 kbp) operon can be routinely inserted in a single copy into *E. coli* strains having one, two or three IS elements within their genomes, relying solely on recombineering. We have no reason to think that IS elements could not be used as recombination targets if they were present at higher copy numbers on the chromosome, just the localization of the transgene insertion becomes more tedious. The absolute efficiencies of obtaining true co-integrants are shown on **Table S2**.

**Figure 3.**
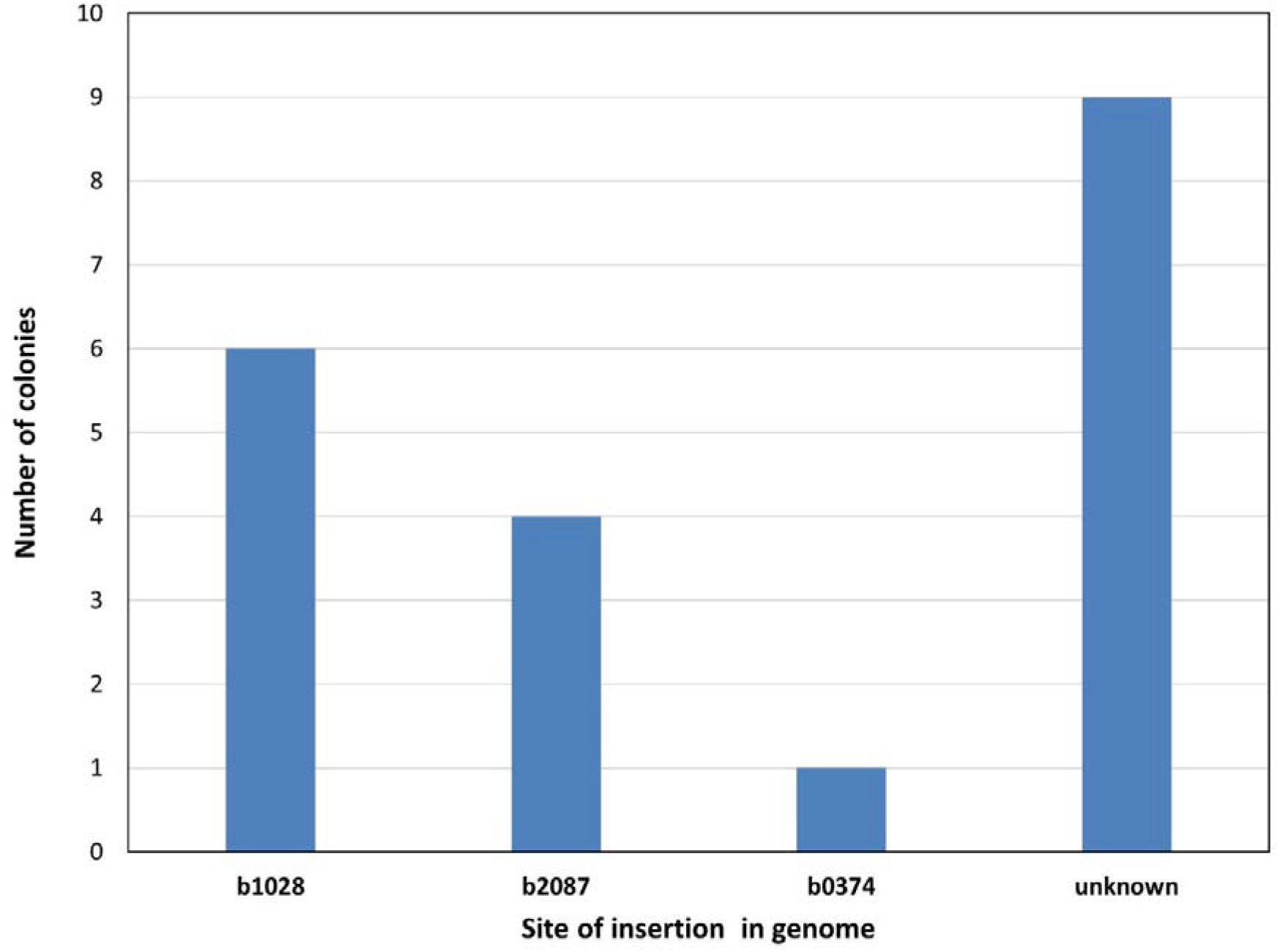
Result of recombineering experiments integrating the VioABCDE_KmR cassette into a chromosomal IS*3* of *E. coli* MDS16. The number of colonies harboring a PCR-verified integration at either of three loci (b1028, b2087 or b03741) or displaying no PCR-verified integration (“unknown”) are shown.

### Targeting ISes by CRISPR-mediated recombineering

After demonstrating the utility of recombineering to load genomic ISes with selectable transgenes, we continued with testing the process of CRISPR/Cas-mediated recombineering of transgenes into genomic IS elements. Among our objectives were to increase both the efficiency of recombineering and the maximum size of the insertable payload as well as to attempt the insertion of unselectable genes and the insertion of a transgene in multiple copies. Our CRISPR/Cas-mediated experiments could be divided into two strategies: on the one hand, we used subsequent CRISPR/Cas cleavage after the recombineering step as an additional tool of counterselection against the wild type (wt) genotype. In these experiments, the λ-Red recombinase enzymes and the CRISPR/Cas machinery were provided by the pORTMAGE2 and pCas9 plasmids, respectively. On the other hand, we also tested concomitant CRISPR/Cas cleavage during the recombineering step to aid the recombination process by generating free DNA ends (in addition to the selective effect mentioned previously). In this latter case, we used the NO-SCAR system (i.e. plasmids pKDsg and pCas9cr4), developed earlier by the Prather laboratory for a similar purpose [27].

When targeting a single copy of IS*1* or IS*3* in E coli MDS42IS1 or MDS30, respectively, subsequent CRISPR/Cas cleavage (with pCas9IS1 or pCas9IS3) was occasionally able to increase the absolute efficiency of recombination, but this effect was inconsistent, with certain cases even falling short of the recombination rates seen in simple recombineering (**Figure S4**: 2 cases improve, 2 cases worsen SpR integration into MDS42IS1). However, when it came to targeting strains harboring two copies of the respective IS element, the advantage of subsequent CRISPR/Cas cleavage became readily apparent: double co-integrants were routinely obtained using multiple protocols. In the end, we simplified the process to two transformations (described in the Methods): the linear DNA is transformed first for recombineering, and the appropriate pCas9 plasmid cleaving the wt form of the IS element is transformed the next day to enforce selection. The pCas9IS1-mediated counterselection in MDS39R2 resulted in 1/7 (14%) or 3/10 (30%) of the tested colonies harboring the resistance cassette at both loci of IS*1* (for short and long homologies, respectively). Counterselection using pCas9IS3 was even better with 15/20 (75%) of the colonies proving to harbor double co-integrants in BLK16. Double IS3 co-integrants were also successfully obtained in MDS27, displaying the portable nature of this technique. **Figure 4** shows the fraction of colonies harboring double co-integrants after one or two rounds of recombineering, using either short or long homologies in MDS39R2.

**Figure 4.**
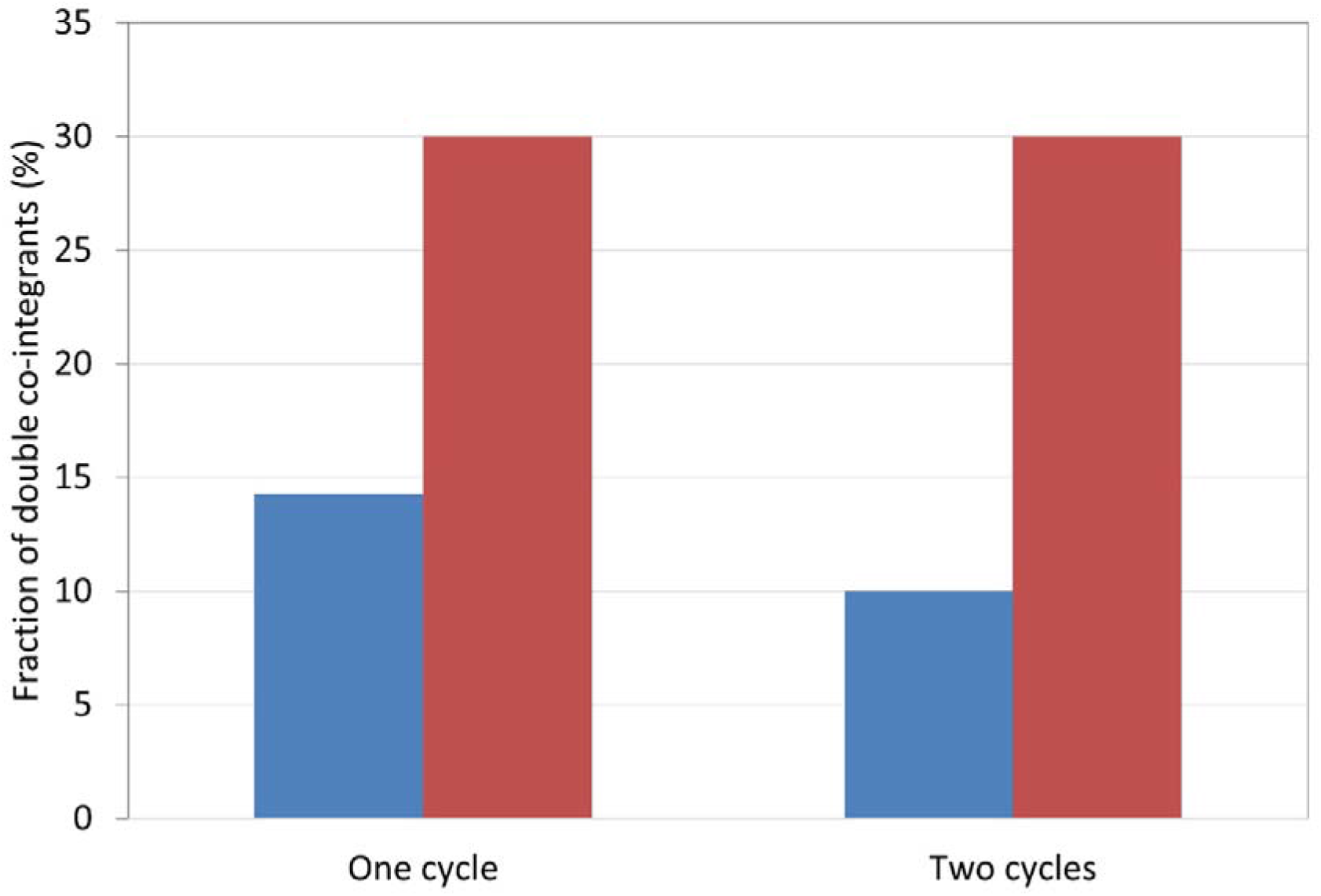
Efficiency of engineering double co-integrants. The IS*1* elements of *E. coli* MDS39R2 were targeted by recombineering followed by consecutive pCas9IS1-mediated counterselection. The fraction of colonies harboring an SpR cassette within both chromosomal IS*1* elements are shown after one or two cycles of recombineering and one round of counterselection. Blue bars depict the use of short (40 bp), red bars depict the use of long (100 bp) homologies.

Besides generating double co-integrants, the great value of applying CRISPR/Cas-based facilitation was unveiled upon the integration of unselectable genes into the chromosome. Peculiarly, we managed to insert the *gfp* gene into the IS*3* of *E. coli* MDS30 using recombineering and subsequent CRISPR/Cas selection. In the course of these experiments, we demonstrated that a 20 nt spacer is equal or superior to a 30 nt spacer in pCas9IS3 concerning efficiency, and up to 20% (2/10) of colonies obtained proved to be positive by PCR. The integration of *gfp* into the IS*1* element of MDS42IS1 was also successfully achieved, but using recombineering and concomitant CRISPR/Cas cleavage provided by the NO-SCAR system (plasmids pKDsg-IS1 and pCas9cr4). In the latter case, 3 out of 30 colonies (10%) were PCR-positive at best. The elevated green fluorescence levels of both engineered strains were verified using a microplate reader (**Figure S5A, B**).

Next, we tested the effect of CRISPR/Cas cleavage on the integration efficiency of long (>9 kbp) selectable DNA cassettes. Since integrating our Vio operon into IS*1* targets by recombineering led to the loss of violacein production (see above), we focused on experiments targeting IS*3*. First, we tested the effect of subsequent Cas cleavage using pCas9IS3. We found that in most of the experiments targeting MDS27, BLK09 or BLK16 (using either SpR or KmR as selection markers), subsequent Cas cleavage not only failed to improve the efficiency of pORTMAGE-mediated recombineering, but on many occasions the recombinant cells were missing altogether. In a few cases nevertheless, we did manage to obtain single VioABCDE-KmR insertions into either of the IS*3* elements of BLK09 (**Figure 5** **and S6**) or BLK16 but double co-integrants were not obtained this way. Second, we tested the effect of concomitant Cas cleavage using the NO-SCAR system. We started by inserting the VioABCDE-KmR cassette into the single IS*3* of MDS30 using NO-SCAR (plasmids pKDsg-IS3 and pCas9cr4). We obtained single co-integrants with a low absolute efficiency (on the order of 10^−2^ correct colonies/ng), the high ratio of correct colonies (6/10) nevertheless permitted the easy detection of true recombinants. The same system was used to target the two copies of IS*3* residing in MDS27, with 7.7% of the colonies displaying the operon inserted at both loci. The absolute efficiencies of obtaining recombinants using CRISPR/Cas-assisted recombineering are listed in **Table S3**.

**Figure 5.**
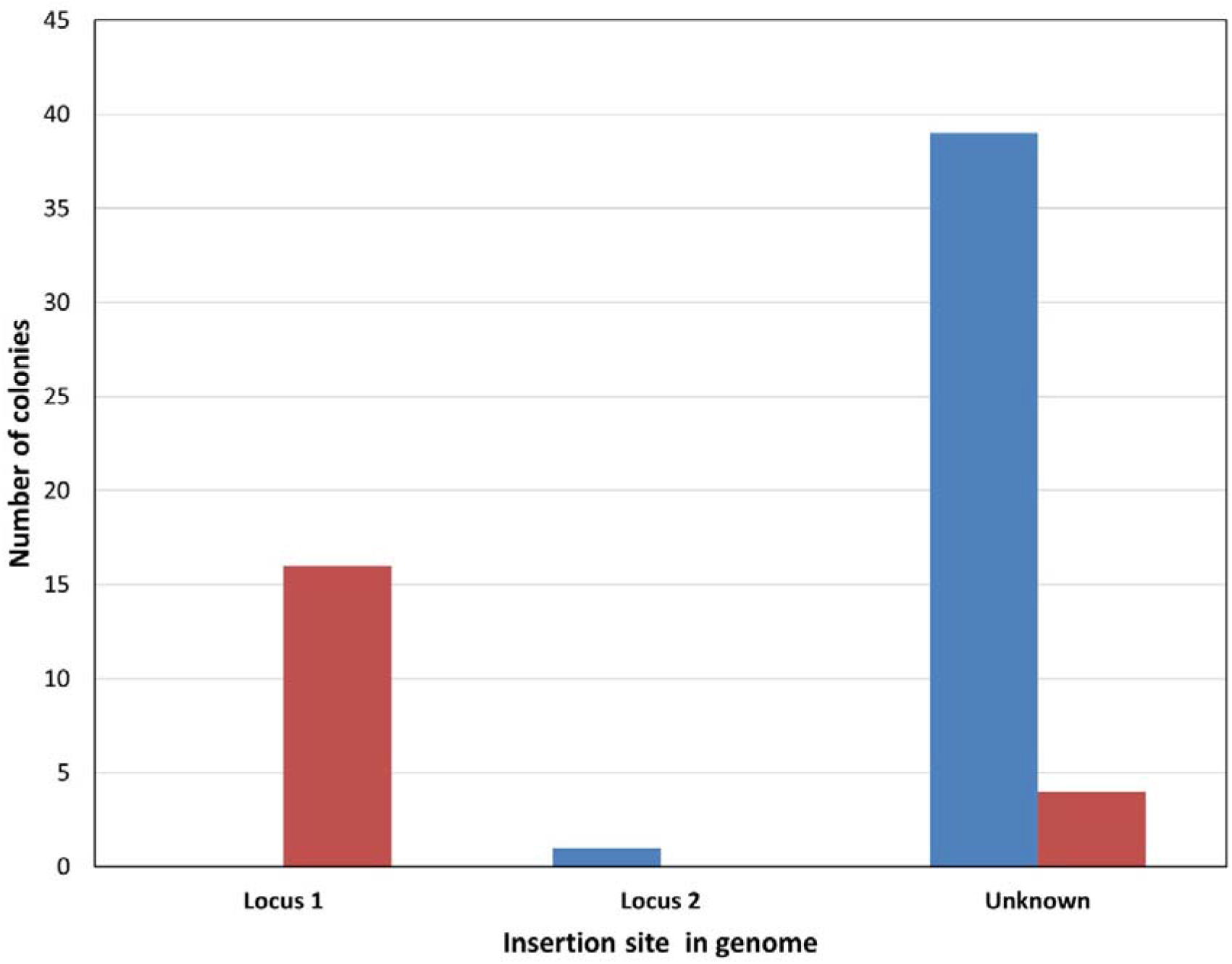
Integration of VioABCDE_KmR cassettes into chromosomal IS*3* elements using recombineering followed by pCas9IS3-mediated counterselection. The number of colonies harboring a PCR-verified integration at locus 1, locus 2, or displaying no PCR-verified integration (“unknown”) are shown for *E. coli* BLK09 (blue) and *E. coli* BLK16 (red). The absolute recombination efficiencies were 4 and 30 recombinants/ng DNA for BLK09 and BLK16, respectively.

### Amplification of cargo genes using copy-paste transposition of IS elements

After verifying that genes and large operons can be integrated into IS elements, our next goal was to achieve the copy-number amplification of these co-integrants via the process of copy/paste transposition. Our initial experiments dealt with the amplification of single resistance genes (e.g. SpR) inserted into IS*1* or IS*3*. The respective transposase genes were expressed from a plasmid in trans, and host cells carrying copy-number amplified resistance genes were selected on antibiotic-gradient plates. Colonies picked from the high-end of the gradient plates went through genomic DNA preparation and ddPCR to quantify the copy number of the resistance gene. These pilot experiments revealed two important conclusions. First, both IS*1* and IS*3* could be used to yield highly-resistant colonies upon transposase induction (**Figures S7AB and S8**). IS*1* experiments however, tended to be less reproducible, with unknown factors influencing the observable increase in resistance (**Figure S7CD**). Second, ddPCR of the initial amplification experiments indicated a major increase in copy numbers of the resistance gene, rising to 20 copies/cell in the first round (**Figure S9, left**). However, when repeating the ddPCR on the same cells after curing them from the transposase-expressing plasmids, the detectable copy numbers fell to the range of 2-4 (**Figure S9, right**). This indicated that most of the high resistance was caused by transposition of the resistance cassette into the transposase-expressing plasmid. We therefore engineered temperature-sensitive transposase plasmids (pSTinsAB’tetR and pSTtnp3tetR for IS*1* and IS*3*, respectively), and plated the cells on gradient plates only after the plasmids had been cured from the cells (as described in the Methods section). This guaranteed that only chromosomal co-integrants of the resistance gene are selected after transposase expression.

Due to the low reproducibility of IS*1*-mediated amplification and to the fact that violacein expression was not observable upon IS*1*-targeted integration (described in the previous section), we focused primarily on amplification of the violacein operon (VioABCDE) using IS*3* elements. Strains of *E. coli* BLK16_IS*3*::VioABCDE_SpR, generated by recombineering (see above) were used as starting points. These carried a single copy of the violacein operon inside the IS*3* element residing at locus 1, along with an SpR marker. After the first round of transposase expression, we analyzed five colonies by ddPCR (targeting the resistance marker) from the high-end of the gradient plates and found the mean copy number to be 5.50 (±1.22) (**Figure 6**). Choosing a colony displaying 7.59 copies and re-expressing the IS*3* transposase led to a further significant elevation of the mean copy number within colonies displaying high resistance levels (8.03±1.21, n=4) (**Figure 6**). Again, we chose the colony displaying the highest copy number (9.10) and expressed the IS*3* transposase for the third time. The mean copy number of the colonies analyzed in the third round did not significantly differ from those of the second round (**Figure 6**). However, by ‘cherry picking’, we were still able to find individual clones displaying further elevated copy numbers (see below).

**Figure 6.**
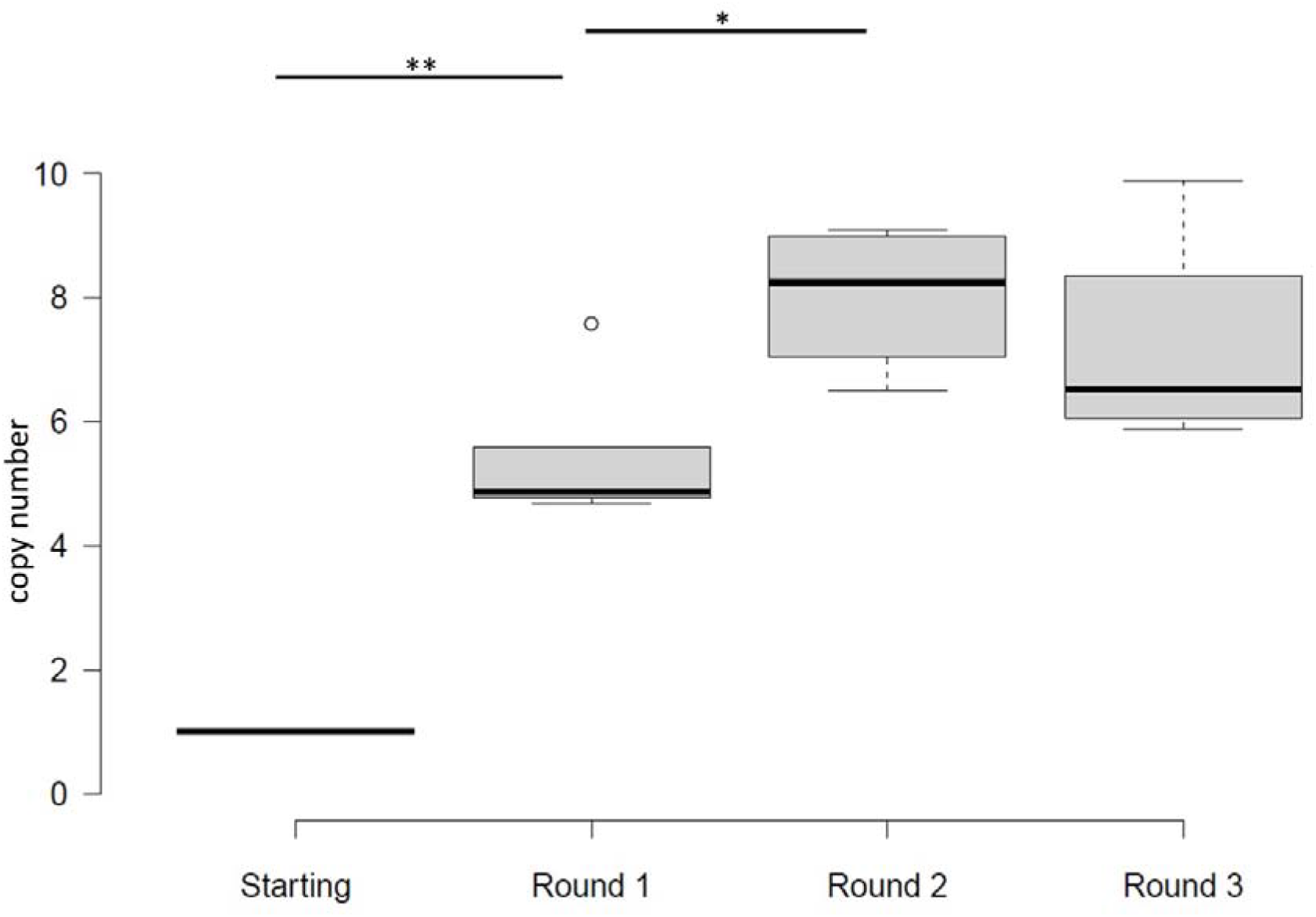
Copy numbers of the SpR gene of *E. coli* BLK16_IS*3*::VioABCDE_SpR detected by ddPCR after 1, 2 or 3 rounds of IS*3* transposase induction. Center lines show the medians; box limits indicate the 25th and 75th percentiles as determined by R software; whiskers extend 1.5 times the interquartile range from the 25th and 75th percentiles, outliers are represented by dots. n = 1, 5, 4, 4 sample points. * p<.05 with two tailed, unpaired t-test; ** p<.002 with two tailed, one sample t-test.

Next, we sent a series of four strains obtained from the above experiment (the starting strain and one clone from each round of transposase induction) for whole genome sequencing. Our aims were i) to verify the copy numbers of the amplified IS*3*::VioABCDE_SpR elements, ii) to analyze the fraction of mutant or truncated VioABCDE operons, iii) to test whether amplification leads to tandem repeats or a random scatter of the loaded IS*3* element within the genome and iv) to check for unexpected genomic rearrangements. The sequence of the starting strain (B0) confirmed the presence of the VioABCDE operon, along with the SpR marker within the IRs of the IS*3* element residing at locus 1 (genomic coordinates 649,839-659,074). The sequence of strain B1, obtained in the first round of transposase induction displayed a 5.71-fold elevated relative coverage of the loaded IS*3*, and unexpectedly, of a 14 kbp segment directly downstream of the IS (amplified region: 649,839-673,907). This stands in good agreement with the 5.59 copy numbers measured by ddPCR. Sequencing of strain B2, which is not a direct descendant of B1, revealed a 11.32-fold increased relative coverage of a similarly large genomic segment delimited on the left side by the left IR of the loaded IS3 (coordinates: 649839-677086). Interestingly, it also displayed a further 5.44 copies (totaling to 16.76) of an interior segment (659074-667565), which is directly downstream of the manipulated IS. The approx. 11-fold increase in the copy number of the violacein operon roughly supports the 9.1 copies indicated by the ddPCR. Finally, strain B3, a descendant of B2, was expected to harbor 26.11 copies of the resistance marker, based on ddPCR. Sequencing did not confirm this, the relative coverage of the genomic region encompassing the loaded IS*3* (649839-677086) nevertheless increased from 11.32 to 12.51, and the number of additional copies of the interior segment downstream of the IS (659074-667565) decreased from 5.44 to 3.54. We therefore concluded that although the third round of induction did not increase the mean copy number of the inserted operon, screening a low number of colonies (<10) permitted the identification of a clone that displayed further amplification of the transgenes. We note that in another clone of the same series of induced strains we detected 9.89 copies, and in another induction series done in parallel, we measured 10.71 and 11.83 copies of SpR using ddPCR. We also experienced that in a small fraction of analyses (1-5%), the ddPCR results were misleading (data not shown), underlining the importance of result verification. The sequenced strains did not harbor the loaded IS*3* scattered throughout the genome, but rather amplified in tandem repeats. However, the IS copies did not form a back-to-back array since, rather unexpectedly, they were amplified together with a large (14-18 kbp) DNA segment lying directly downstream to them.

Violacein content of the starting strain and its derivatives obtained after various rounds of transposase induction were routinely measured. The amount of violacein released from the BLK16IS*3*::VioABCDE_SpR derivatives displayed a good correlation (R^2^=0.98) with the copy number of the VioABCDE operon inferred from its relative sequencing coverage (**Figure S10**). Copy numbers of the SpR marker determined by ddPCR, relative violacein content and the relative sequence coverage of the violacein operon (acquired by whole genome sequencing) are displayed on **Table 3** for each sequenced strain of the IS*3*::VioABCDE_SpR-amplification process done in BLK16.

**Table 3.** Properties of *E. coli* BLK16IS*3*::VioABCDE_SpR derivatives after the indicated rounds of IS3 transposase induction

| Strain code | Strain name | SpR copy number by ddPCR | Violacein content (AU) | Relative violacein content<sup>1</sup> | Relative sequence coverage of VioABCDE by WGS<sup>2</sup> |
| --- | --- | --- | --- | --- | --- |
| B0 | BLK16IS3::VioABCDE_SpR, starting | 0.98 | 0.06 | 1 | 1 |
| B1 | BLK16IS3::VioABCDE_SpR, 1st round | 5.59 | 0.2926 | 4.88 | 5.71 |
| B2 | BLK16IS3::VioABCDE_SpR, 2nd round | 9.10 | 0.7847 | 13.08 | 11.32 |
| B3 | BLK16IS3::VioABCDE_SpR, 3rd round | 26.11 | 0.8113 | 13.52 | 12.51 |

SpR: Spectinomycin resistance
ddPCR: droplet digital polymerase chain reaction
AU: arbitrary unit, based on OD<sub>585</sub> measurement
WGS: whole genome sequencing
<sup>1</sup> Relative to strain B0
<sup>2</sup> Ratio of sequence reads obtained +/- 10 nt of peak border

To verify the general applicability of the inPOSE strategy, we tested the amplification using IS*3* elements residing elsewhere in the same strain, employing further strains of *E. coli*, and also using a different selection marker. Inducing the IS*3*::VioABCDE_SpR at locus 2 in strain BLK16 for one round also yielded purple colonies with significantly elevated copy numbers of the operon (3.28±0.15) (**Figure S11**). Importantly, when analyzing white colonies by ddPCR, the mean copy number was even higher (8.20±2.79). In this experiment, we induced a white colony for two further rounds, and identified white colonies harboring ever higher copy numbers of the SpR marker (31.70 and 49.14 in round 2 and 3, respectively), the increase of the means, however were not significant (**Figure S11**). One round of transposase induction also worked in BLK09IS3::VioABCDE_SpR (locus 2) (**Figure 7**), the obtained mean copy number was 3.88 (±1.68). Interestingly, using KmR in the same locus, we only found a maximum copy number of 2.27, with the mean not being significantly elevated (1.51±0.52).

**Figure 7.**
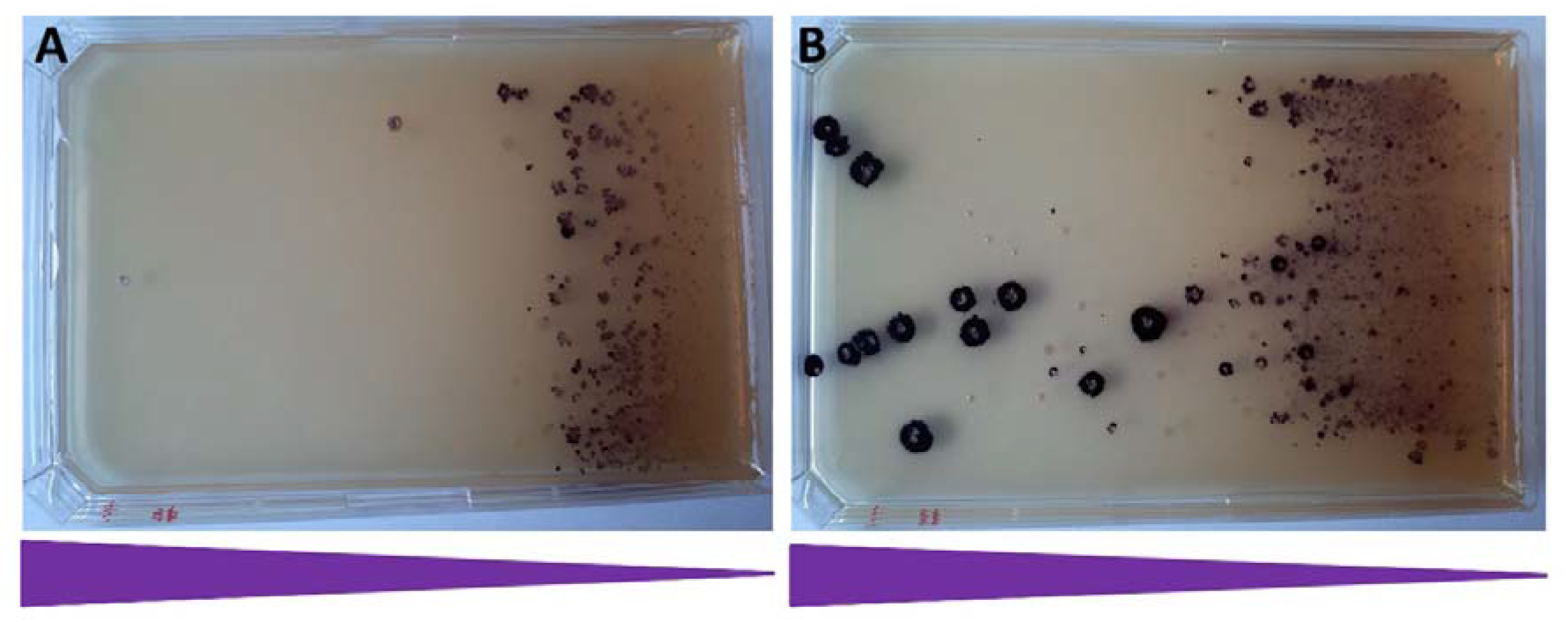
The effect of one round of IS*3* transposase induction on the Sp-resistance of BLK09IS*3*::VioABCDE_SpR (locus 2) colonies. Note the intense purple colors of the colonies caused by violacein. (A) Uninduced cells, (B) cells induced with aTc. The triangles represent the gradient of Sp concentration within the medium.

To test a completely different genetic background, amplification was also tested in strains MDS27 and MDS30, derivatives of *E. coli* K-12. For *E. coli* MDS30IS*3*::VioABCDE_KmR, one round of transposase induction boosted the number of highly resistant colonies compared to the uninduced (**Figure S12**). The mean copy number of the KmR marker was also significantly elevated (7.57±2.39), however, only a small fraction of the colonies growing on the plate were purple. For *E. coli* MDS27doubleIS3::VioABCDE_KmR, a strain that had both copies of IS*3* loaded with VioABCDE_KmR at the starting point, we carried out three rounds of induction. Similarly to that seen for MDS30IS3::VioABCDE_KmR, the first round of induction elevated the copies of KmR marker to 4.78 (±0.76), but most of the colonies were white (**Figure S13**). The second and third rounds of induction, both of which were carried out choosing a purple colony, could not significantly increase the mean copy number of the marker (5.7±2.66 and 4.0±0.64, respectively). Again by cherry picking, we could find colonies carrying up to 7.8 copies. Importantly, the second and third rounds of this experiment revealed that in the lack of induction, the majority of the colonies remain purple (**Figure S14** and **S15**), but the obtainable mean copy numbers are somewhat limited (4.07±1.96 and 2.95±1.16, respectively).

### Stability of genomic co-integrants

A key phenotypic trait of engineered strains is the stability of expressing the inserted transgenes in the lack of selection. We compared strains harboring chromosomal VioABCDE operons to those carrying the same operon on multi-copy plasmids using two different tests. Our first test monitored the fraction of cells displaying violacein production. We chose three well known strains, *E. coli* BL21(DE3), DH5α, and MG1655 to be our controls, all transformed with the pUTLIQ_vioABCDE plasmid. From our collection of genome-engineered strains we tested three versions of BLK16IS3::VioABCDE_SpR (carrying 1, 5.59 or 10.5 copies of IS*3*::VioABCDE_SpR, respectively), and strain MDS27IS*3*::VioABCDE_KmR, carrying 5.8 copies, according to our ddPCR measurements. As apparent on **Figure 8**, all three control strains displayed a steep decrease in the ratio of purple colonies, and completely lost the purple phenotype by generation 40. On the contrary, all four genomic co-inegrants tested in this experiment retained purple color in >96% of their colonies, with the single copy strain not displaying loss of function at all. We note that i) multi-deletion strains (MDS42 and BLK16) of *E. coli* displayed complete plasmid loss even faster than their conventional, non-reduced counterparts, and ii) further single and the double copy-harboring strains [BLK16IS*3*::VioABCDE_SpR (1 copy, locus 2) and MDS27IS*3*::VioABCDE_KmR, 2 copies, respectively] did not display any loss of function either (**Figure S16**). We therefore conclude that our genomic co-integrants can be used for the prolonged expression of transgene arrays in the lack of selection with a negligible fraction of cells displaying complete loss of production.

**Figure 8.**
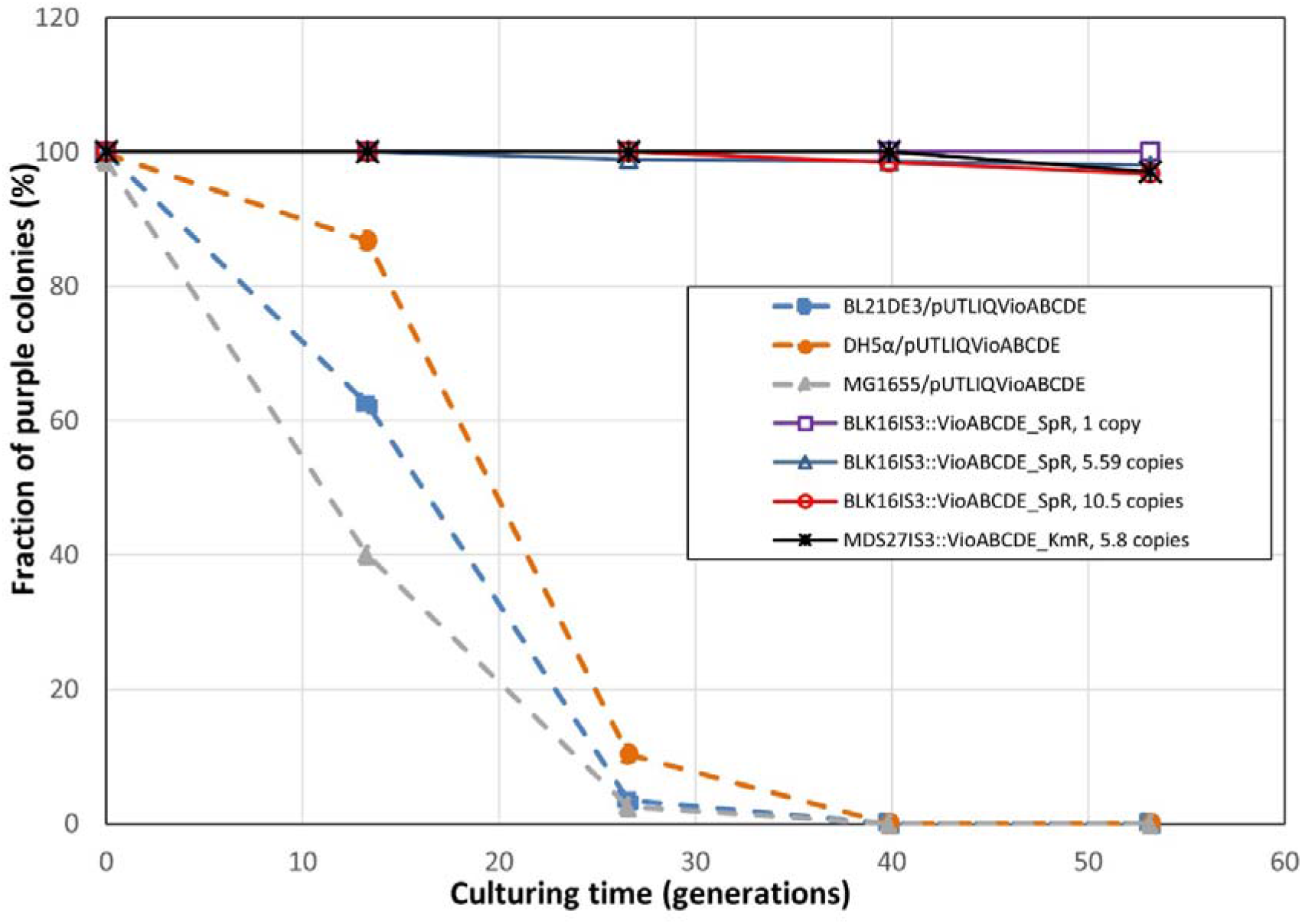
Fractions of violacein-producing cells within bacterial cultures grown in the lack of antibiotic selection. Dashed lines mark strains carrying the pUTLIQvio_ABCDE plasmid, solid lines mark strains carrying an IS*3*::vioABCDE cassette on their chromosomes, as indicated on the legend. All values are means of three biological replicates.

Our second test assaying the stability of violacein production in the lack of selection measured the levels of violacein extracted from the cultures each day. As expected, we saw the rapid decline of violacein levels in case of *E. coli* BL21(DE3), MG1655 and MDS42 strains initially harboring the pUTLIQ_vioABCDE plasmid (**Figure 9**). *E. coli* DH5α displayed a somewhat smaller, but still significant loss of 35% (P=0.012 with a two tailed, unpaired t-test). In contrast, no significant decrease in the violacein levels were observed on day 4 (compared to day 1) for strains BLK16IS*3*::VioABCDE_SpR carrying 5.59, 9.1 and 10.5 copies of the violacein operon, respectively, and strains MDS27IS*3*::VioABCDE_KmR carrying 2 and 5.8 copies, respectively. Furthermore, in this test the violacein levels produced by MDS27IS*3*::VioABCDE_KmR carrying 5.8 copies significantly surpassed that of all other strains, including the four strains carrying the pUTLIQ_vioABCDE plasmid (**Figure 9, Figure S17**).

**Figure 9.**
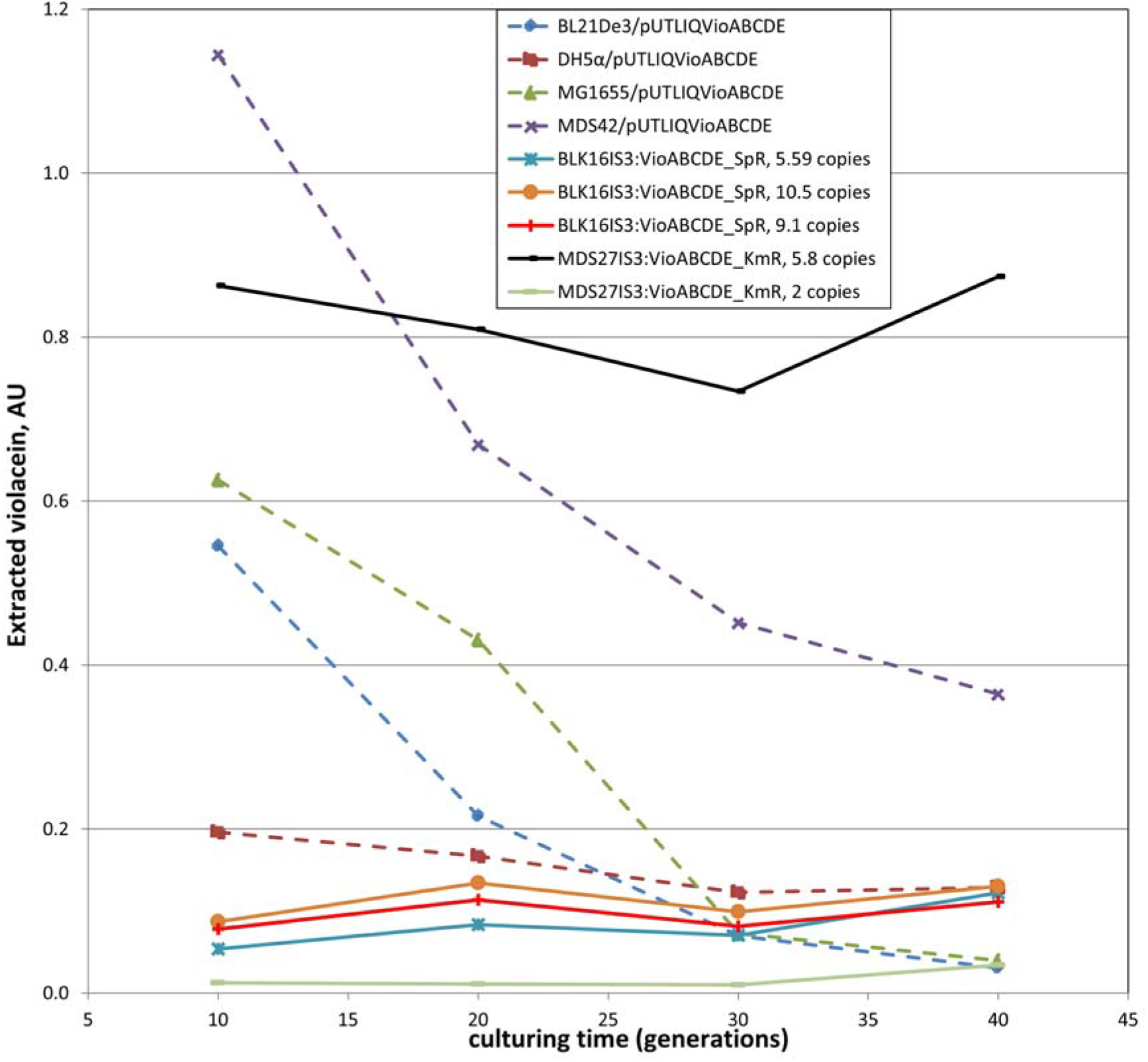
Violacein levels of liquid bacterial cultures grown in the lack of antibiotic selection. Dashed lines mark strains carrying the pUTLIQ_vioABCDE plasmid, solid lines mark strains carrying an IS3::vioABCDE cassette on their chromosomes, as indicated on the legend. All values are means of three biological replicates.

### Transposition of marked IS*3* elements into the *E. coli* chromosome

Finally, we asked whether IS*3* can be transferred into a bacterial strain of choice by controlled transposition. This process, which can be referred to as a potential “Step 0”of the inPOSE protocol (**Figure 10**) could be valuable for future users who wish to utilize IS*3* for chromosomal cloning and amplification inside a strain that has no copies of this element at all. For the process of IS*3* entry into the chromosome, we decided to use transposition, catalyzed by the plasmid-encoded transposase used in the transgene copy-amplification experiments above. To make the IS*3* entry selectable, a marked version of IS*3* was required, we therefore used the IS*3*::SpR allele generated in the course of SpR recombineering in MDS30 (see section on “Targeting ISes by recombineering” above). The IS3::SpR cassette was PCR-amplified from the chromosome and cloned in to a suicide plasmid that is unable to replicate in strains lacking the *pir* gene. This donor plasmid, pSG78A_full_IS*3*::SpR was transformed into *E. coli* MDS42 carrying the transposase-expressing pSTtnp3tetR plasmid, followed by plating on Sp plates. Replica plating revealed that 10% (2/20) of the obtained colonies were Sp^R^Cm^S^Ap^S^, indicating that they had lost both the donor plasmid and the transposase plasmid, but retained the SpR gene, presumably due to its transposition into the genome. To verify transposition and identify the locus of entry, we carried out ST-PCR on the chromosomal DNA prepared from two colonies. Two pairs of PCR reactions were carried out, as described in the Methods (**Figure S18**). Sequencing a unique product (from SmFwd + CEKG4 PCR on colony 11) confirmed IS*3*::SpR insertion into the *pqiA* gene at position 1,012,519 (using NC000913.3 coordinates). In a similar experiment, the transposition of IS*3*::SpR into MDS30 was induced and selected, and this time, 33% (10/30) of the colonies displayed a Sp^R^Cm^S^Ap^S^ phenotype. Most of the ST-PCR reactions carried out on the genomic DNA of ten such colonies generated unique PCR patterns, probably indicating different points of insertion (**Figure S18**). Sequencing a PCR product from one of the least complex patterns (Sp347 fwd + CEKG4 PCR product of colony 8) revealed IS*3*::SpR integration into the 5’untranslated region of the waaL gene (position 3,796,945, NC000913.3 coordinates). We therefore demonstrated the transposition-mediated entry of IS*3*::SpR into two different strains of *E. coli* (one IS-free, the other harboring a single copy of IS*3*), with IS*3*::SpR entering the chromosome at two distinct loci.

**Figure 10.**
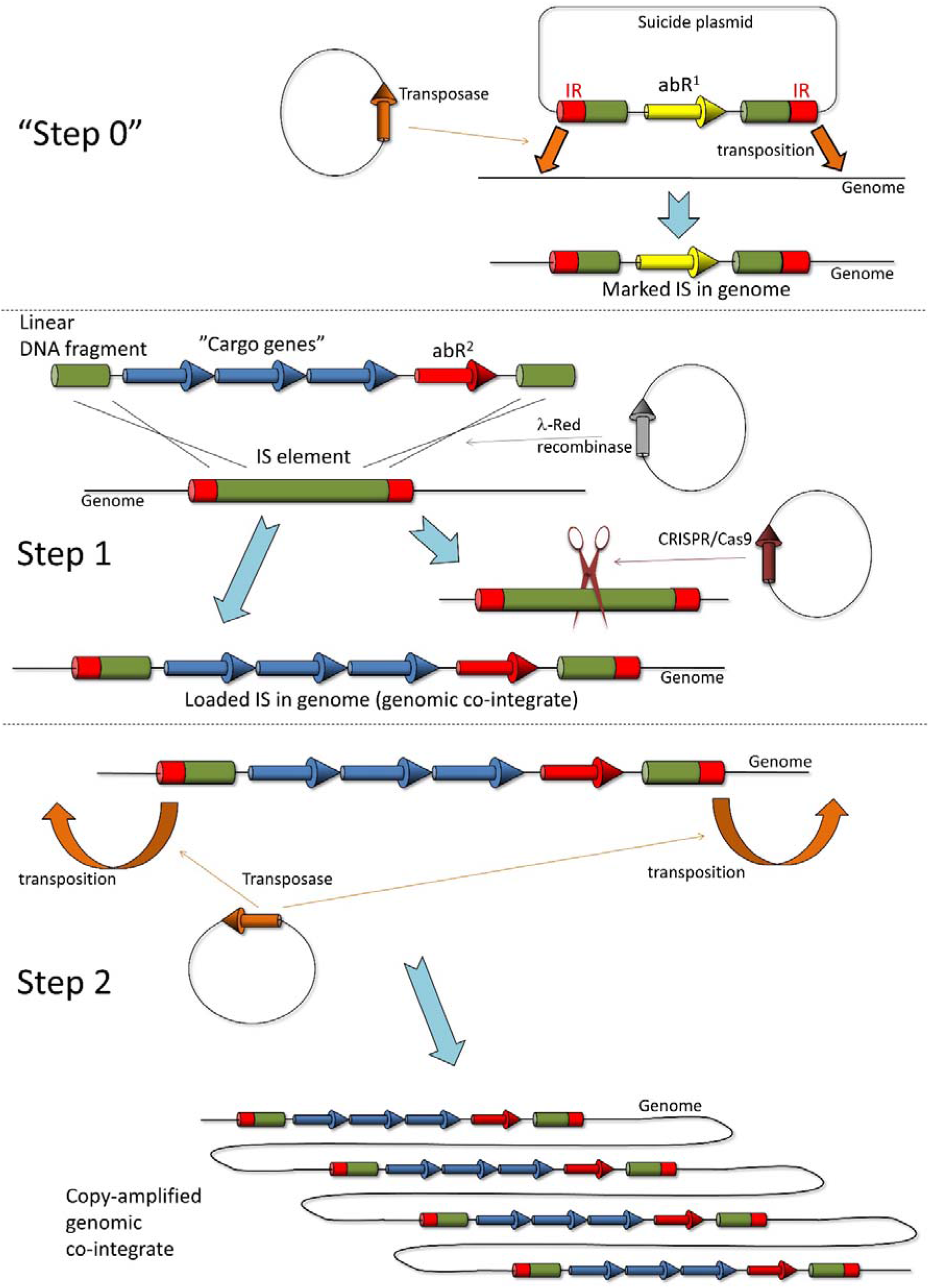
The inPOSE protocol. Step 1 is the entry of a gene or operon of interest into an IS element residing on the host genome by recombineering, facilitated by the λ-Red recombinase enzymes. CRISPR/Cas-mediated cleavage of the wt IS element(s) enforces selection for the recombinants and (if applied concomitantly) facilitation of the recombination event. In Step 2, the genomic co-integrant (i.e. the loaded IS) is copy-amplified by the transposase corresponding to the IS, expressed in trans. “Step 0” is an optional accessory step that can transpose the marked IS element into the genome of host cells chosen for chromosomal transgene cloning. Green boxes: the targeted IS element (or segments thereof); red boxes: inverted repeats (IRs) of the IS; yellow and red arrows: two different antibiotic resistance genes (abR^1^ and abR^2^, respectively); blue arrows: the transgenes to be integrated; orange arrows: transposase gene of the targeted IS; gray arrow: λ-Red recombinase genes; brown arrow: CRISPR/Cas genes; open thin black lines: host bacterial genome; closed thin black lines: circular plasmids.

## Discussion

The interest towards techniques capable of rapidly integrating genetic constructs into the genome has been present for decades. The first such methods relied on homologous recombination of plasmids with the chromosome [73]. Others used phage integrases and chromosomal attachment sites for site-specific integration of the plasmid [4, 74]. This technique was later optimized to allow integration of plasmid assembly mixes instead of finished plasmids, to permit dual integration in a single step, and to remove the selection marker from the co-integrate warranting scalability of integration [75]. A further strategy exploited the FRT site-specific recombinase target site and its respective recombinase, FLP to introduce plasmids into the chromosome [76]. This method required the prior insertion of the acceptor FRT sites into the genome by random Tn*5* transposition. The high target-selectivity of Tn*7* was also utilized to insert genetic cargo at the Tn*7* chromosomal attachment site by transposition from a donor plasmid [77, 78]. Our method is related to most previously described techniques, yet it is different: it uses mobile elements as landing pads, i.e. recombination target sites, instead of utilizing their transposase activity for integration. The recombination process is mediated by the λ-Red recombinase enzymes expressed from a plasmid, providing portability to our method. The transposase activity is nevertheless used, but only in the second stage of our process: the copy-number amplification of the inserted transgene(s).

The recombineering of single selectable transgenes into IS elements worked as expected. The achieved absolute recombination efficiencies were mostly in the range of 0.5-20 recombinants/ng DNA, which meets or exceeds efficiency values reported in the literature [24, 79], and are sufficient to conveniently obtain co-integrants in a single try. If one wishes to generate chromosomal libraries, the use of longer homologies could be a viable solution; in our experiments, increasing homology length 2.5-fold (to 100 bp) elevated recombination efficiency over 13-fold to up to 268 recombinants/ng DNA (**Table S2**). The positive surprise was brought about by our recombineering experiments dealing with >9 kbp-long DNA fragments. Despite our low expectations based on earlier reports [24], we readily obtained single co-integrants of the marked VioABCDE operon in the chromosomes of strains carrying one, two or three copies of IS*3*. Although our PCR screens also suggested formation of single co-integrants when targeting IS*1* elements, the lack of VioABCDE function prevented us to use these strains further. We suspect a technical fault during the generation of the linear fragments as the reason; this however was not investigated further. When inserting a single resistance gene into strains with a single copy of the targeted IS, all obtained colonies carried the transgene within that IS. When two IS*3* copies were present in the genome, still all integrations took place at known loci. In all other cases however, a certain fraction of the obtained colonies did not harbor the transgene(s) in any of the IS copies. We called these events integrations at unknown loci, which happened more often when targeting IS*1* or when attempting to insert a long DNA cassette, possibly indicating a higher rate of illegitimate recombination.

Next, we complemented recombineering with CRISPR/Cas cleavage, which we expected to confer four advantageous effects: i) increase the rate of recombination by providing free DNA ends, if used concomitantly [12, 65]; ii) increase the maximum size of the payload that is insertable [24]; iii) allow the insertion of unselectable genes and iv) allow insertion of transgenes in multiple copies in a single step. The use of CRISPR/Cas to aid selection of recombinants or to facilitate the recombination process did not result in the increase of recombination efficiency when targeting a single IS in the genome. However, obtaining double co-integrants was only possible with CRISPR/Cas selection. This is not surprising, for as long as an unloaded (wt) IS remains in the genome, the Cas9 ribonucleoprotein will find and cleave it, and introducing a double-strand break into the chromosome is a very powerful form of negative selection in *E. coli*. We did not manage to obtain triple co-integrants however, when targeting the three copies of IS*3* found in MDS16. This could indicate that the frequency of triple recombination is lower than the frequency of double strand break-repair, allowing the formation only of double recombinants. Perhaps transforming the linear DNA in multiple (≥3) cycles prior to CRISPR/Cas selection could solve this issue, this however was not attempted. Besides obtaining double co-integrants, the use of CRISPR/Cas was also essential in the process of recombining an otherwise unselectable gene (*gfp*) into the chromosome. Despite our attempts, we were not able to obtain double *gfp* integration into the genome possibly indicating that applying multiple mechanisms of selection (i.e. resistance and Cas cleavage) has a synergistic effect on the formation of recombinants. The use of CRISPR/Cas cleavage concomitantly or subsequently to the recombineering step did not show noteworthy differences when inserting single genes into the genome. However, integrating the >9 kbp DNA cassette in multiple copies into the *E. coli* chromosome was only possible using the NO-SCAR system. We attribute this to the facilitating effect of free DNA ends (generated by the concomitant cleavage) on recombination efficiency.

The next milestone of our project was the amplification of the inserted cargo. Although multiple strategies are available in the literature to integrate transgenes into the bacterial genome, very few projects have dealt with the multi-copy cloning of heterologous genes or operons in the bacterial chromosome. The first example used the transposase of Tn*1545* to generate a library of insertions, marked with an antibiotic resistance. In a second step, the inserted cassettes were accumulated in one strain using P1 transduction, always removing the selective marker after each transfer using the *xis* and *int* functions of phage λ [80]. Another work used a phage integrase to insert the DNA cassette and the FLP/FRT system to eliminate the selection marker, thereby allowing further integrations at different phage attachment sites [75]. These two projects managed to accumulate the *lacZ* gene in the *E. coli* chromosome in three and four copies, respectively. The third example differs from the two prior since it did not use serial re-integration, but applied true gene copy number amplification. The co-integrate was marked with an antibiotic resistance gene, and serially passaging cells in increasing antibiotic concentration allowed selection for spontaneous RecA-mediated gene duplications arising from unequal crossovers between the daughter chromosomes [81]. Copy numbers up to ~40 were achieved using that strategy, which was ended by deleting the *recA* gene of the host cell to stabilize the tandem repeats. Our strategy is related to the latter one, it however uses the plasmid-expressed transposase of the targeted IS for the amplification process. It does not require an increasing selection pressure to evolve the multi-copy strain, the elevated antibiotic levels present on a gradient plate are required only for the “readout” of the transposition process.

In our gene-amplification experiments, the induction of the transposase had a marked effect on the distribution of resistance levels within the population in the first round (**Figure S7, S8**). In the second and third rounds of induction however, the difference in the distribution of resistance levels between the induced and uninduced population was smaller, if noticeable (**Figures S14, S15**). This may indicate that at higher starting copy numbers of the marked IS, even the leaking expression of the transposase was sufficient to facilitate enough transpositions to generate highly-resistant cells. As an alternative explanation, the spontaneous RecA-mediated duplication of marked ISes most likely takes place at all copy numbers; the duplication of many copies (at later rounds) however probably has a more dramatic effect on resistance than the duplication of a single copy (in the first round). Either way, these spontaneously arising clones of elevated resistance, even if arising less frequently, can harbor just as many copies as the induced clones.

A key parameter of transgene-carrying strains is the stability of transgene expression, especially in the absence of antibiotic selection. Our first experiment measuring this parameter declared colonies as producers and non-producers in a binary fashion, based on their purple or white color, respectively. This experiment indicated a rapid loss of the producer phenotype for plasmid-carrying strains, but the maintenance of production by >96% of the cells in the case of genomic co-integrants (**Figure 8**). In a more quantitative analysis, we measured the violacein levels extractable from liquid cultures grown in the lack of selection. Again, the significant decrease of violacein levels was seen only in the case of plasmid-based expression, and not in the case of chromosomal expression (**Figure 9**). The strong strain dependence of violacein production was also apparent in the second type of experiment, both for plasmid-based and chromosomal transgene expression.

The utility of the inPOSE system for transgene entry by homologous recombination, and transgene amplification by copy/paste transposition was demonstrated in the experiments above. We have no reason to think that IS*3* is the only element applicable for this purpose, nor is it necessarily the best. In fact, we only refer to inPOSE as a strategy, not a discrete tool, and believe that other mobile elements should be tested for this purpose to be applied in further bacterial strains or species. For those future users, who wish to use IS*3* for this purpose, but their targeted bacterial cell lacks this element, we showcase a transposition-based method to insert a marked copy of IS*3* into the genome of the chosen host bacterium. We refer to this as “Step 0” of the inPOSE protocol, which can be readily continued with Step 1 if using a different antibiotic for transgene insertion.

Despite the multiple advantages of using inPOSE for chromosomal gene cloning and amplification, several shortcomings of the system are yet present. First, it seems that not all copy-and-paste IS elements will be useful for this strategy, as apparent from the suboptimal performance of IS*1*-mediated integration and amplification in the course of our work. We currently do not possess sufficient information to explain the behavior of IS*1*, therefore we cannot predict which further IS elements will be optimal choices for this type of task. The systematic testing of further ISes should provide us more insight concerning the nature of the problem. Second, the transposition of the loaded ISes did not result in a random scattering of the IS within the chromosome, but in the formation of tandem repeats instead, along with the amplification of the 14-18 kbp-long genomic segment lying directly downstream of the IS. This is probably the result of the transposase enzyme not acting on the right IR of the IS, but instead on a pseudo-terminus, lying further away. Transposases acting on pseudo-termini of transposons have been described [82, 83], the question here is why the transposase misses the canonical right terminus of the loaded IS*3*. The reason may be that the 827 bp missing from the center of the interrupted IS*3* removes an unknown protein or RNA factor necessary for correct transposition, and the transposase gene expressed in trans fails to complement this missing function due to its codon-optimized nature. We will try to identify this factor and provide it *in vivo* to obtain controlled transposition that is equivalent to the natural process. And third, the expression levels of the integrated transgenes and operons is in certain cases unexpected, or hard to predict. A notorious example can be seen on **Figure 9**: MDS27IS*3*::VioABCDE_KmR strains carrying 5.8 copies of the KmR gene (measured by ddPCR of the KmR gene) produces much more violacein than the BLK16IS*3*::VioABCDE_SpR strain carrying 9.1 or 10.5 copies. On the contrary, BLK16IS*3*::VioABCDE_SpR strains displayed violacein production that was more or less proportional to the copy number of the operon, as apparent on **Figure 9**, and **Table 3**. This obviously points to the strain dependence of violacein production, and underlines the general necessity of phenotypical testing, besides analyzing copy numbers by genetic methods. Other possible causes of unexpected production levels are the fact that ddPCR measures the resistance gene, not the presence of the complete, correct operon and one must keep in mind the potential errors of ddPCR mentioned previously. When considering long-term stability of genomic co-integrants, one should also take into account the occasional loss of tandem repeats by homologous recombination. The initiation of this process is probably the cause of the slight decrease seen on **Figure 8** for the multi-copy insert carrying strains after 25-40 generations of growth. We hope that recombination-mediated loss of operon copies can be avoided by re-engineering the transposition process, as mentioned above. Until then, an alternative solution could be the removal of the *recA* gene from the genome. We have not tested yet whether and how our induced transposition performs in *recA*- strains. If transposition is seriously impaired, one can remove the recA gene at the end of the induced transposition process, analogously to the prior study applying transgene amplification [81].

## Conclusions

We have shown successful genome editing of *E. coli* by recombineering targeting IS*1* or IS*3* elements, using CmR, SpR or KmR as selection markers. We have demonstrated integrating solely a resistance gene or a five-gene operon marked with a resistance gene this way. Facilitating the recombineering step with CRISPR/Cas cleavage (applied either concomitantly or subsequently) aided the integration of long (>9 kbp) cassettes and was essential to insert unselectable genes (e.g. *gfp*) or to obtain double co-integrants of selectable gene cassettes in a single step. We have displayed the reproducible and controllable copy-number amplification of IS*3*-carried transgenes by expressing the respective transposase in trans. Our work also demonstrated the increased stability of chromosomal transgenes in the lack of selection, compared to their plasmid-borne counterparts. The flexibility of this strategy, called inPOSE, is showcased using the IS*3* element, but is most likely applicable to other bacterial ISes as well. For those who wish to use IS*3* for this purpose in strains harboring no IS*3* copies, we developed tools to rapidly introduce a marked copy of IS*3* into the chromosome by transposition. We look forward to learning about the experience of other laboratories testing inPOSE for chromosomal gene cloning.

## Acknowledgements

We thank Lajos Pintér for next generation sequencing and bioinformatics. This work was supported by the National Research, Development, and Innovation Office of Hungary (NKFIH) Grant No. K119298, the GINOP-2.3.2-15-2016-00001, and The Biotechnological National Laboratory Grant.

