## Supplementary figures for "inPOSE: a flexible toolbox for chromosomal cloning and amplification of bacterial transgenes"

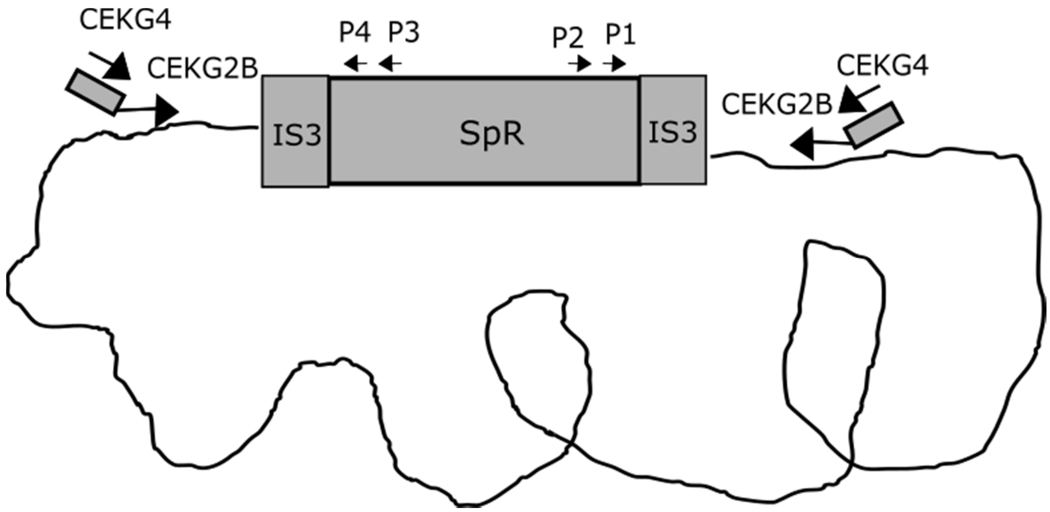


**Figure S1**. Relative positons of primers used for ST-PCR. P2 (Sp439Rev) and P3 (Sp110Fwd) primers were used for 1st PCR, P1 (SmFwd) and P4 (Sp347Fwd) primers were used for 2nd PCR.

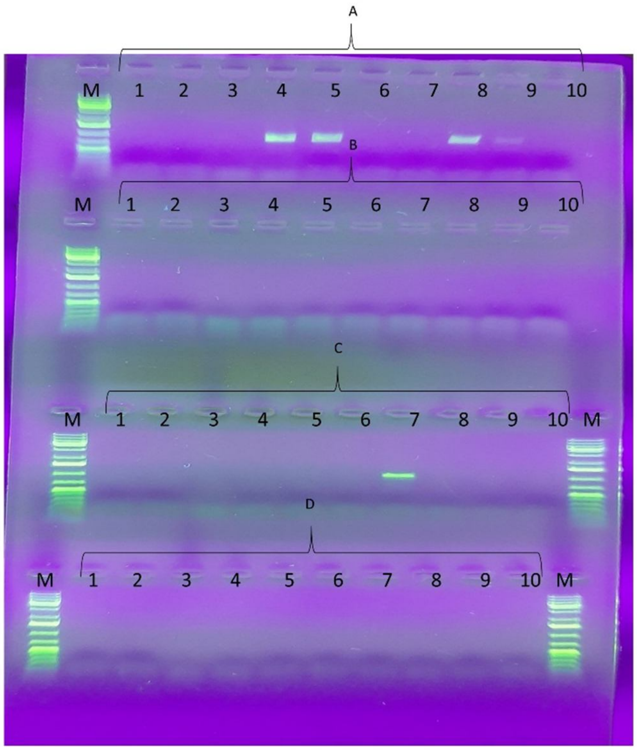


**Figure S2.** Targeting the two genomic IS*1* elements of MDS39R2 with the SpR cassette using pORTMAGE2-mediated recombineering (one of three parallel experiments). Ten colonies obtained upon the recombineering process are screened using four primer sets: A: SmFw + YeaE3, B: SmRev + YeaE3, C: SmFw + aisE2, D: SmRev + aisE2. Four colonies were positive for integration at *yeaJ* locus, and one colony was positive for integration at *ais* locus. M: GeneRuler 1 kb Plus DNA Ladder (Thermo Scientific)

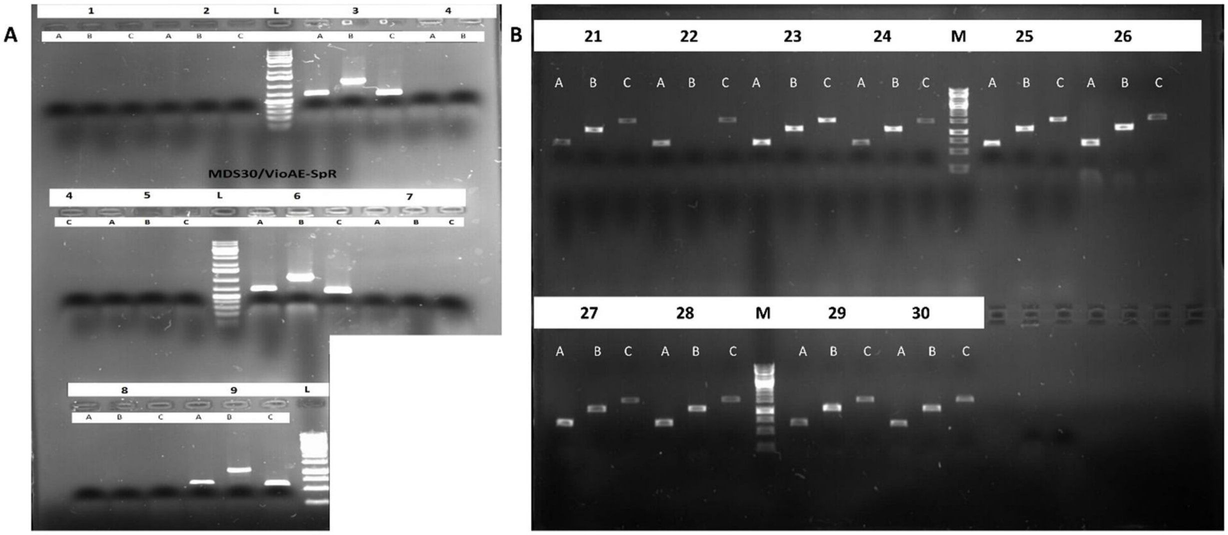


**Figure S3**. The pORTMAGE-mediated insertion of the VioABCDE operon into MDS30. (**A**) Engineering MDS30 with the IS*3*::VioABCDE_SpR cassette. Nine colonies obtained upon the recombineering process are PCR-screened using three primer sets: A: VioDF1 + VioDR3, B: VioAR2 + b1028rev, C: b1025fwd + SmFw. Three colonies (colonies 3, 6 and 9) display positivity. L: GeneRuler 1 kb Plus DNA Ladder (Thermo Scientific) (**B**) Engineering MDS30 with the IS*3*::VioABCDE_KmR cassette. Ten colonies obtained upon the recombineering process are PCR-screened using three primer sets: A: VioDF1 + VioDR3, B: VioAR2 + b1028rev, C: b1025fwd + Km_HindIII_Rev. Successful PCR-amplification with primer pair B shows genomic integration of vioABCDE pathway for colonies 21 and 23-30. M: GeneRuler 1 kb DNA Ladder (Thermo Scientific)

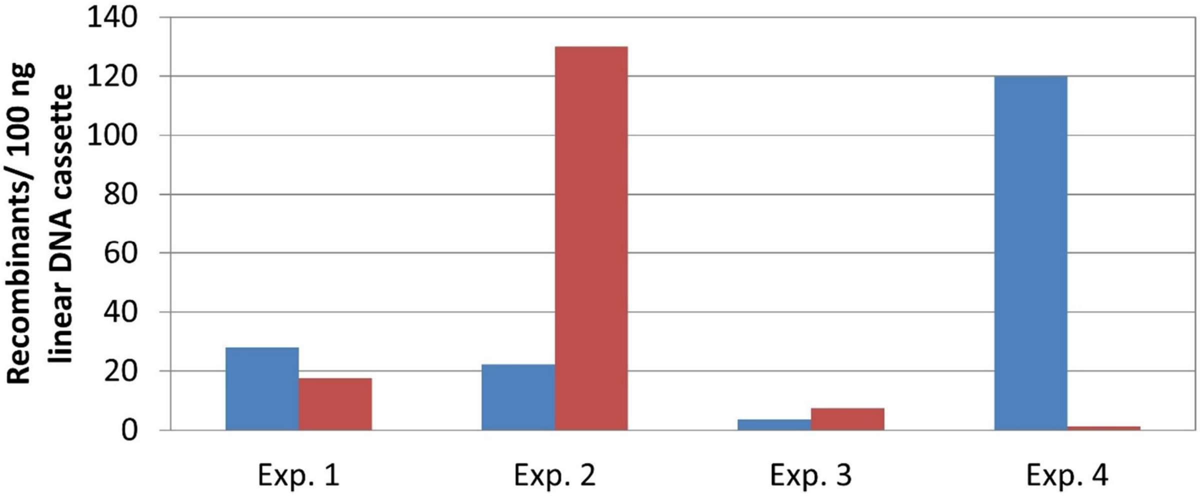


**Figure S4.** Testing the effect of CRISPR/Cas cleavage ont the efficiency of λ-Red recombinase-mediated recombineering into the genomic IS*1* of MDS42IS*1*. The SpR cassette was transformed along with the pCas9IS1 plasmid into target cells harboring induced pORTMAGE2 plasmid. Red bars indicate the efficiency of recombineering alone, blue bars indicate the efficiency of CRISPR/Cas-mediated recombineering.

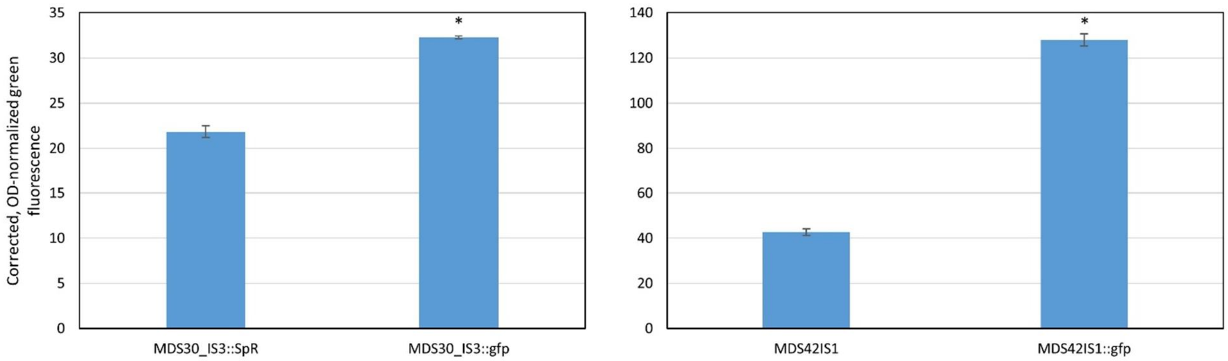


**Figure S5**. Effect of chromosomal *gfp* integration on the fluorescence of the host strain. The fluorescence of fully grown cultures are shown, recorded in a Synergy2 microplate reader (Excitation: 500 nm; Emission: 540 nm). Fluorescence was corrected by subtracting the blanks, and normalized to OD600. The means of three technical replicates are shown. * p<2E-5 with a two-tailed, unpaired t test

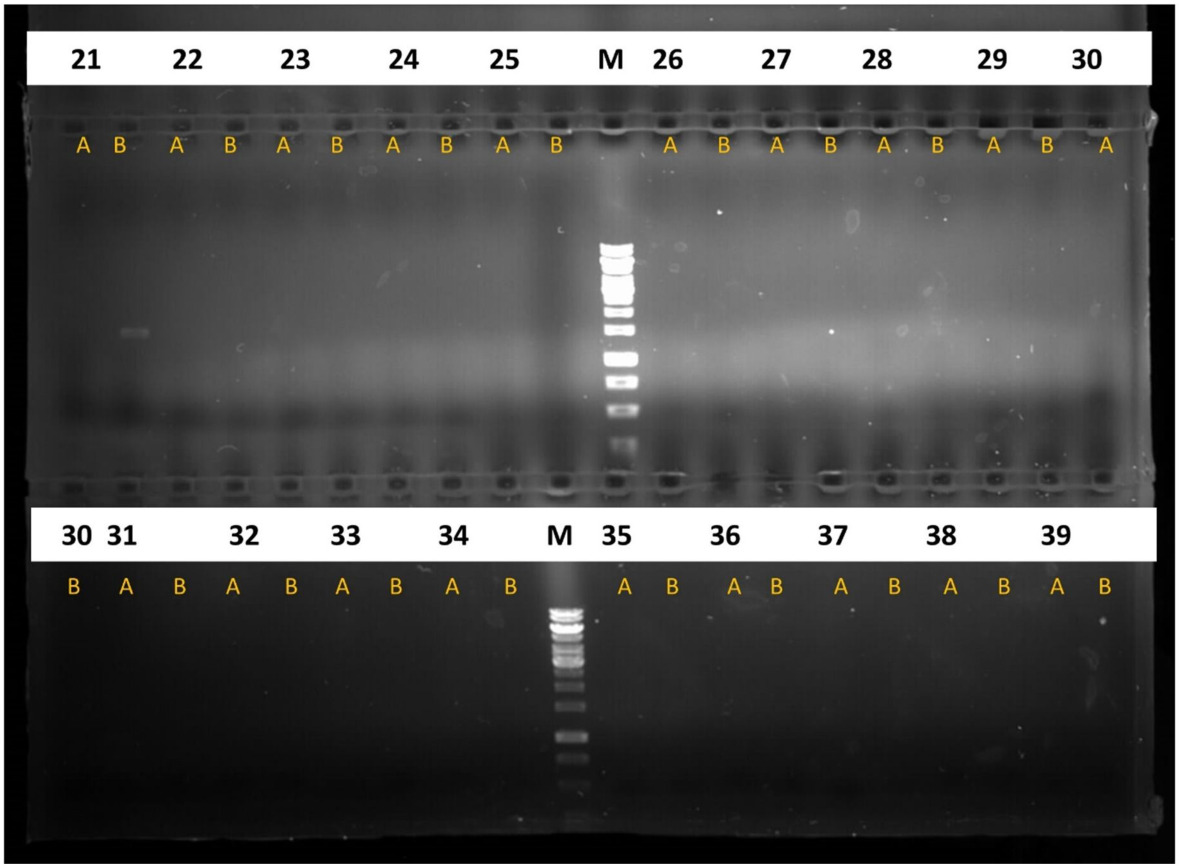


**Figure S6.** Targeting BLK09/pORTMAGE2 cells with IS3::VioABCDE_KmR cassettes, followed by pCas9IS3 transformation. Nineteen colonies obtained upon the CRISPR/Cas-mediated recombineering process are PCR-screened using two primer sets: A: IS3flanking1 + VioAR2, B: IS3flanking2 + VioAR2. One positive colony is detected. M: GeneRuler 1 kb DNA Ladder (Thermo Scientific)

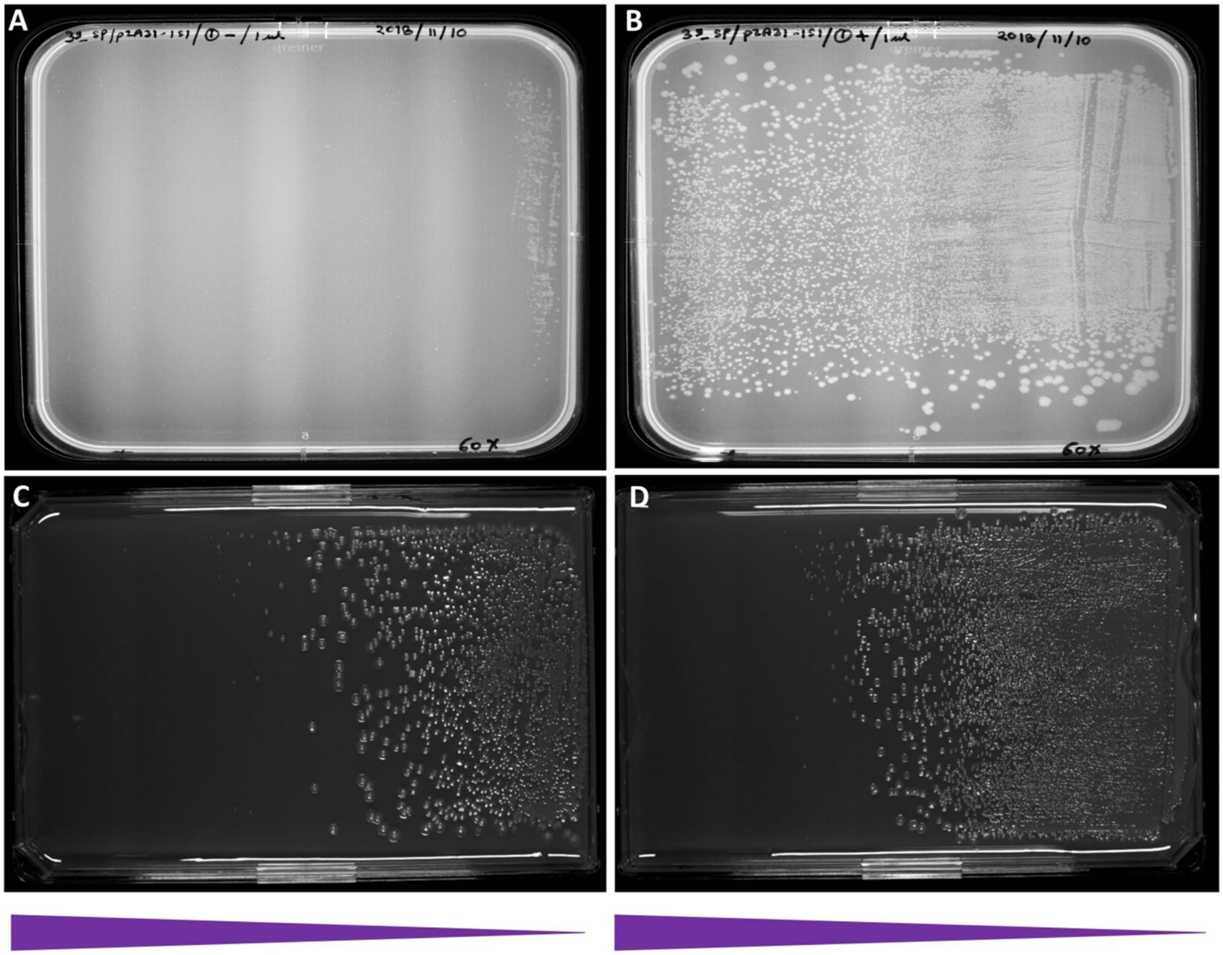


**Figure S7**. The effect of IS*1* transposase induction on Sp-resistance. (A) Uninduced MDS39R2IS*1*::SpR cells, (B) MDS39R2IS*1*::SpR cells induced with aTc. (C) Uninduced MDS42IS*1*::vioABCDE_KmR cells, (D) MDS42IS*1*::vioABCDE_KmR cells induced with aTc. The triangles represent the gradient of Sp concentration (A,B) or Km concentration (C,D) within the medium.

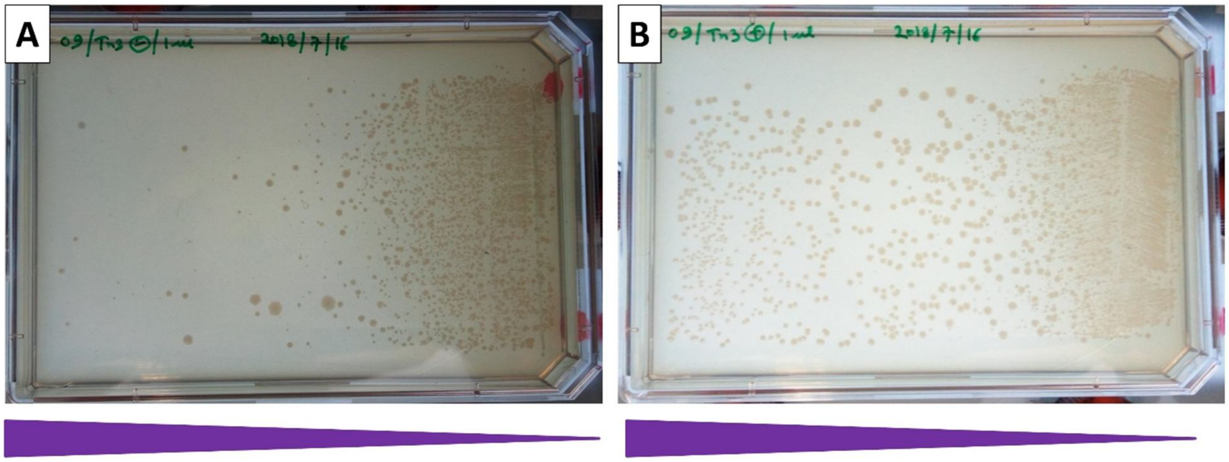

**Figure S8**. The effect of IS*3* transposase induction on the Sp-resistance of BLK09IS*3*::SpR colonies. (A) Uninduced cells, (B) cells induced with aTc. The triangles represent the gradient of Sp concentration within the medium.

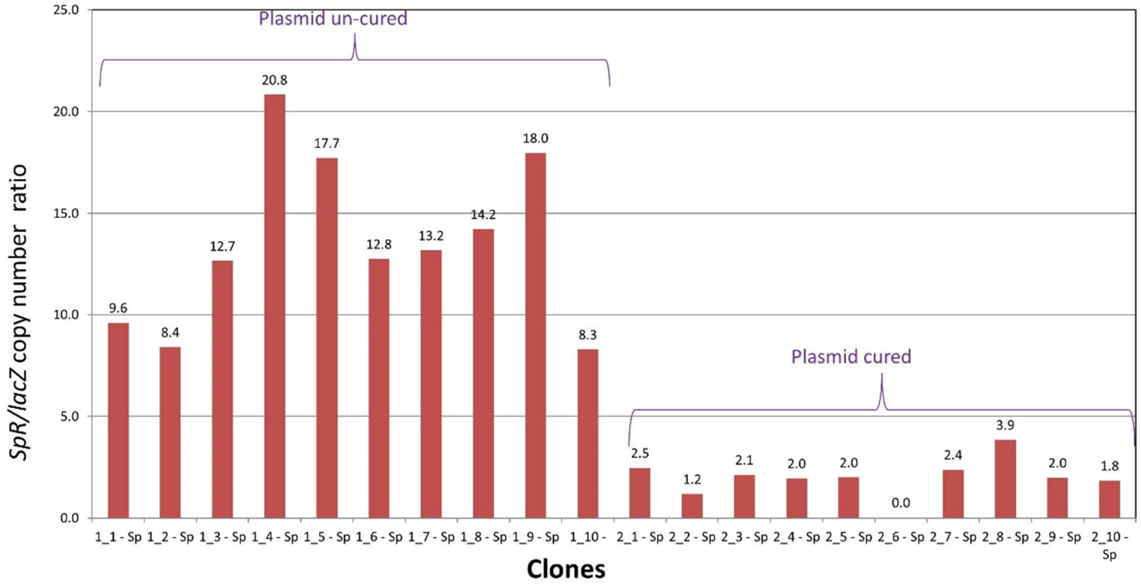


**Figure S9**. Quantification of the *SpR* gene in MDS39R_IS*1*::SpR genome by ddPCR after one round of IS*1* transposase induction. Brackets indicate clones analyzed prior to plasmid curing or after plasmid curing.

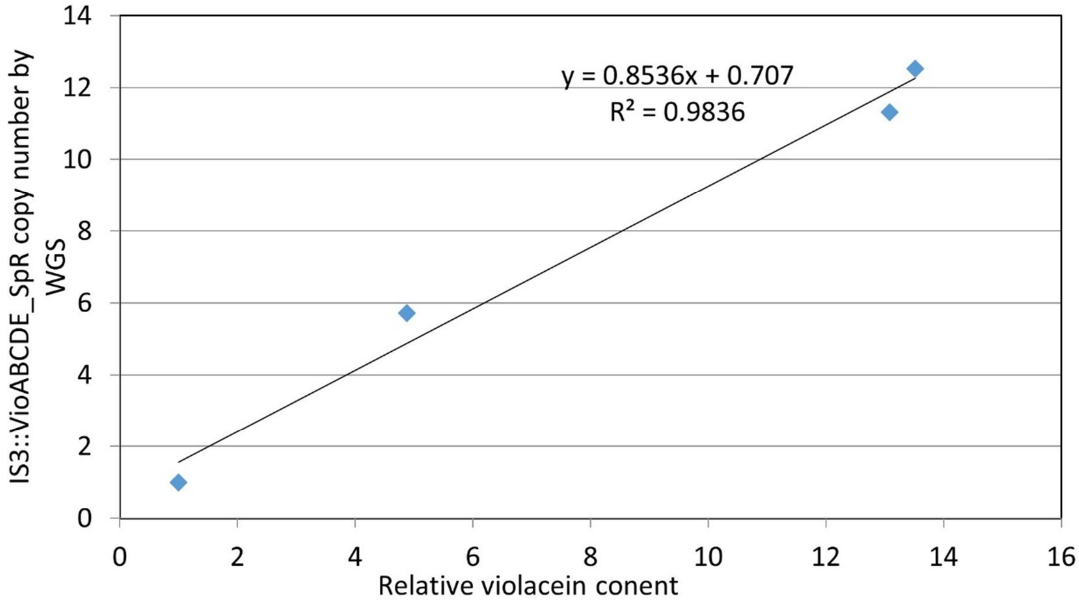


**Figure S10.** The correlation between the relative violacein content of various BLK16_IS*3*::VioABCDE_SpR strains obtained after induction of IS3 transposase, and their corresponding copy number of the genomic IS*3*::VioABCDE_SpR cassette detected by whole genome sequencing (WGS).

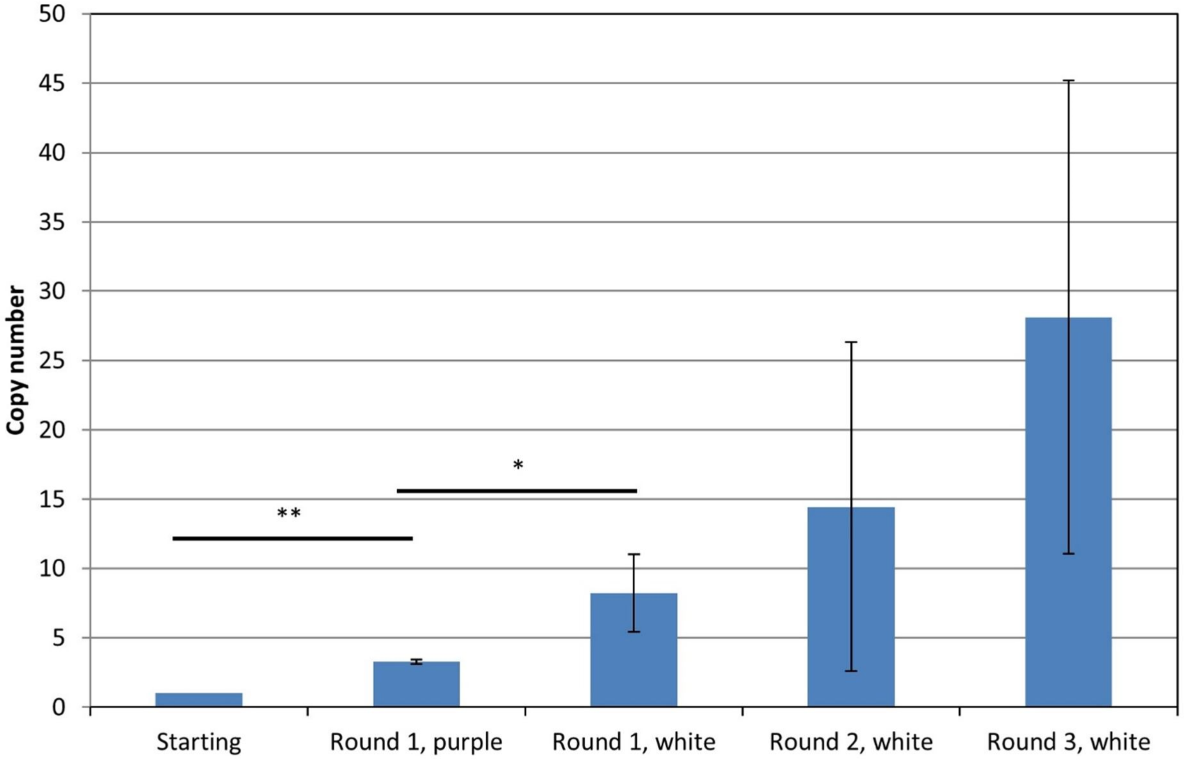


**Figure S11.** Copy numbers of the SpR gene of *E. coli* BLK16_IS*3*::VioABCDE_SpR (at locus 2) detected by ddPCR after 1, 2 or 3 rounds of IS*3* transposase induction. Note that after round 1 of transposase induction, the purple and the white colonies are displayed in separate bars. n = 1, 3, 3, 4,6 sample points. * p<.05 with two tailed, unpaired t-test; ** p<.002 with two tailed, one sample t-test.

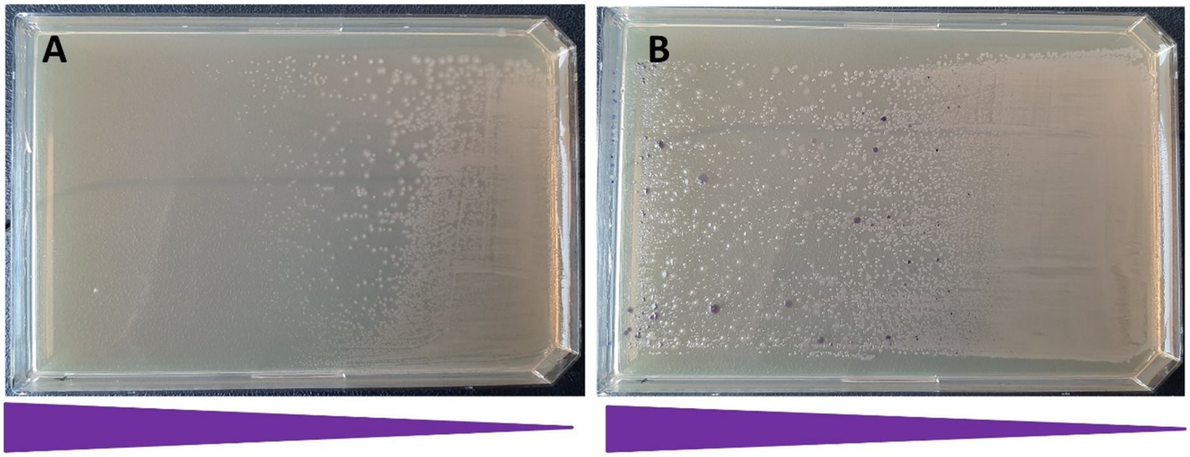


**Figure S12.** The effect of one round of IS*3* transposase induction on the Km-resistance of MDS30_IS*3*::VioABCDE_KmR colonies. (A) Uninduced cells, (B) cells induced with aTc. Note that the majority of colonies are white. The triangles represent the gradient of Km concentration within the medium.

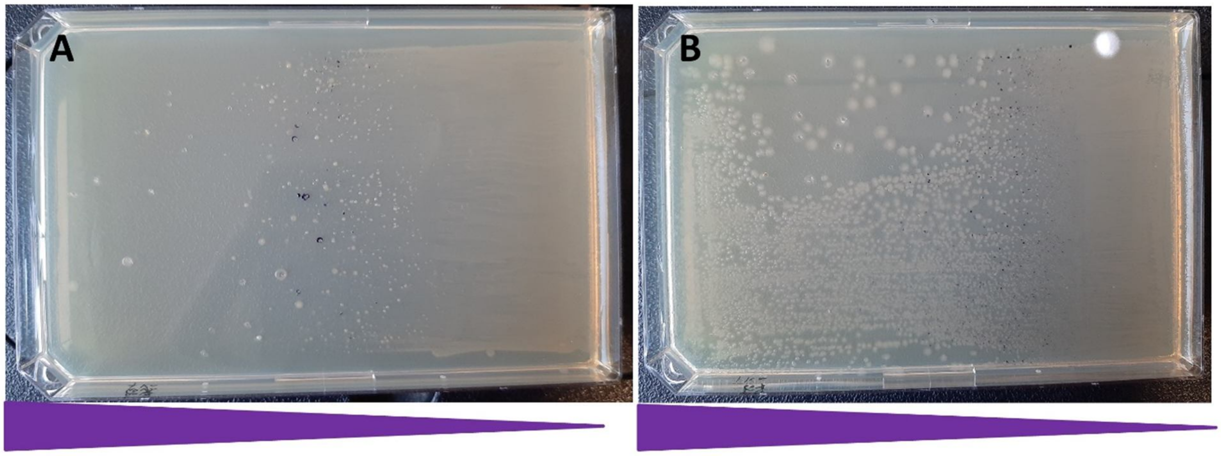


**Figure S13**. The effect of one round of IS*3* transposase induction on the Km-resistance of MDS27_doubleIS*3*::VioABCDE_KmR colonies. (A) Uninduced cells, (B) cells induced with aTc. Note that the majority of colonies are white. The triangles represent the gradient of Km concentration within the medium.

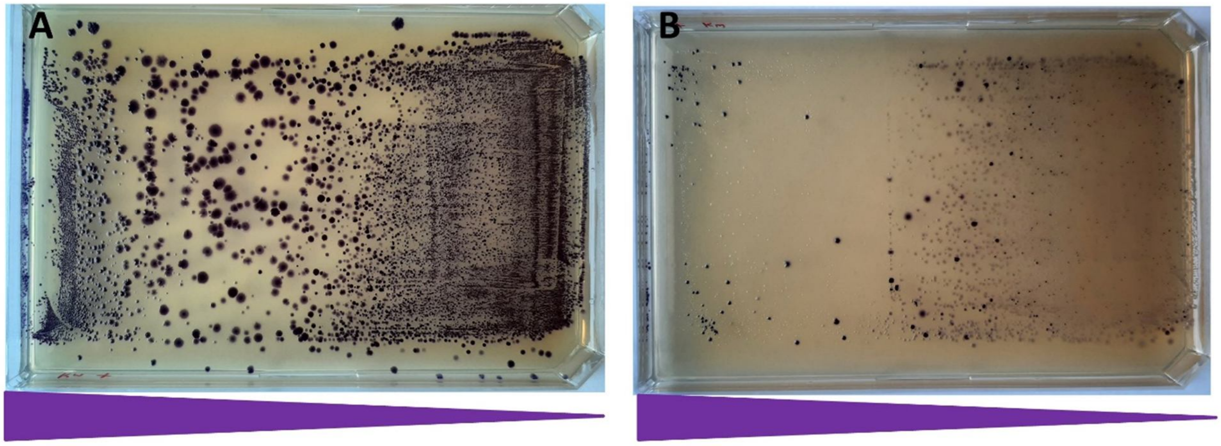


**Figure S14.** The effect of the second round of IS*3* transposase induction on the Km-resistance of MDS27_doubleIS*3*::VioABCDE_KmR colonies (starting copy number: 5.8 copies). (A) Uninduced cells, (B) cells induced with aTc. Note that the majority of induced colonies are white. The triangles represent the gradient of Km concentration within the medium.

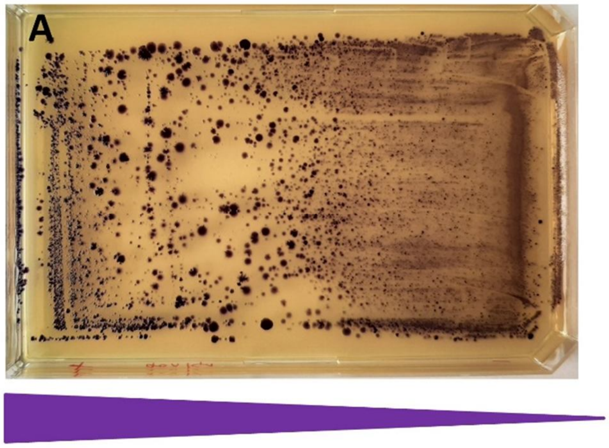


**Figure S15.** The effect of the third round of IS*3* transposase induction on the Km-resistance of MDS27_doubleIS*3*::VioABCDE_KmR colonies (starting copy number: 5.7 copies). (A) Uninduced cells, (B) cells induced with aTc. Note that the majority of induced colonies are white. The triangles represent the gradient of Km concentration within the medium.

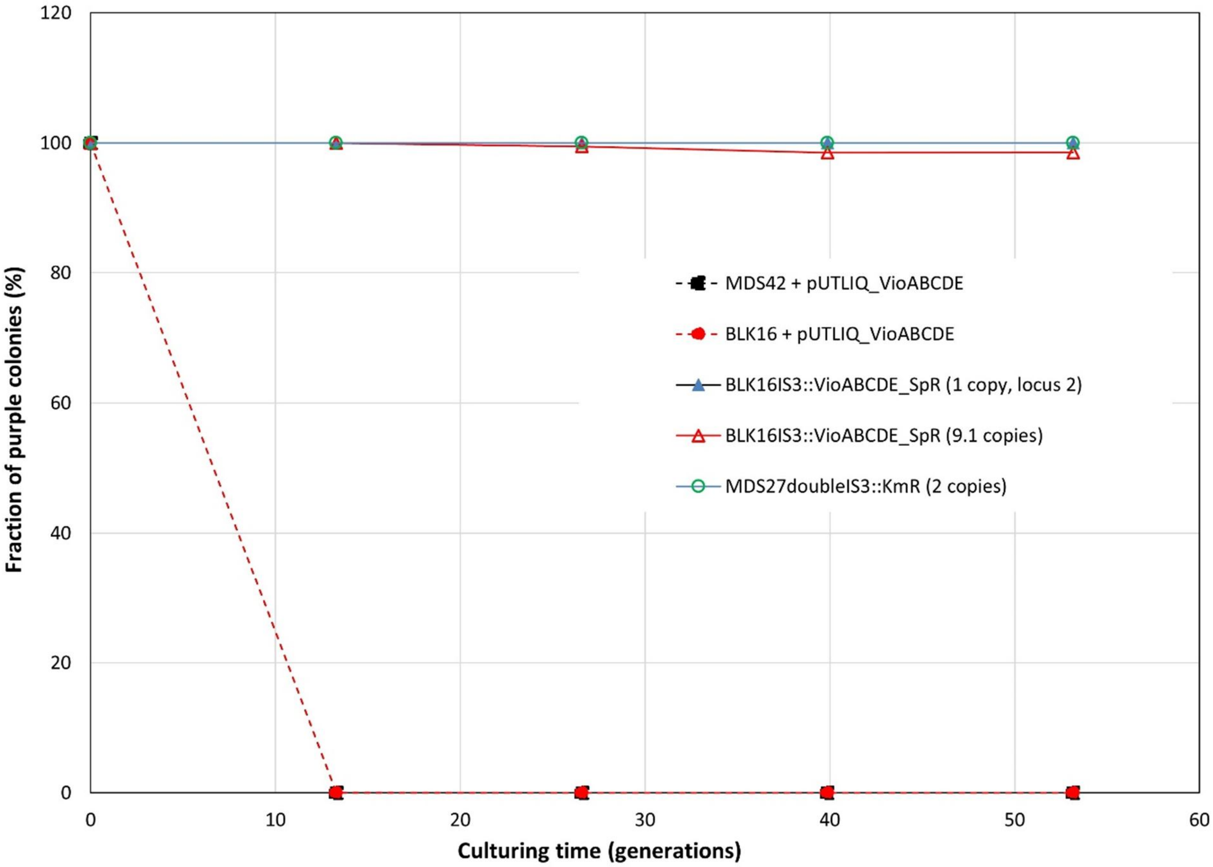


**Figure S16.** Fractions of violacein-producing cells within bacterial cultures grown in the lack of antibiotic selection. Dashed lines mark strains carrying the pUTLIQvio_ABCDE plasmid, solid lines mark strains carrying an IS*3*::vioABCDE cassette on their chromosomes, as indicated on the legend. All values are means of three biological replicates.

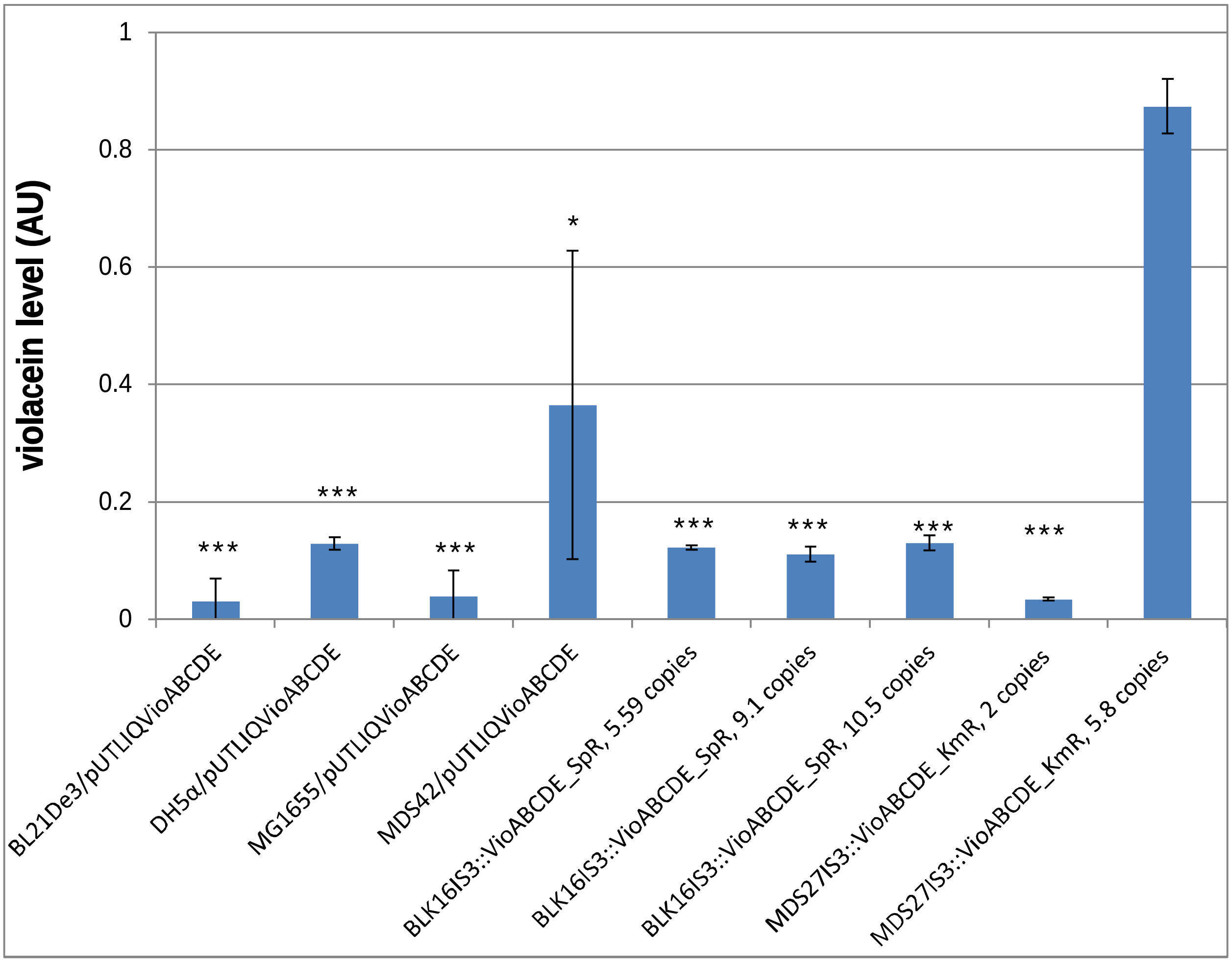


**Figure S17**. Levels of violacein extracted from liquid bacterial cultures after 4 days of growth in the lack of antibiotic selection. Values are means of three biological replicates. Asterisks mark the results of two-tailed, unpaired t-tests comparing the given strain to *E. coli* MDS27IS3:VioABCDE, 5.8 copies (last column) (* $p<0.05$; *** $p<1E-4$).

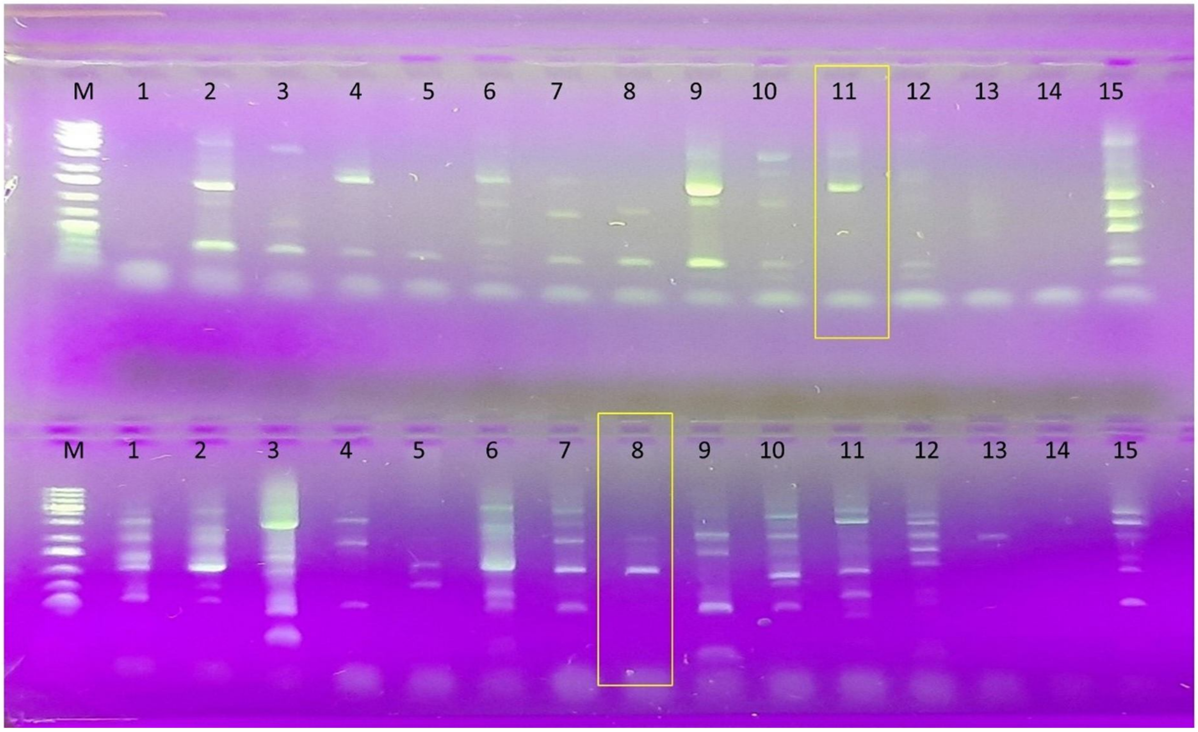


**Figure S18**. Verifying the transposition of IS*3*::SpR from pSG78A_full_IS*3*::SpR by ST-PCR. The figure shows gel electrophoresis of ST-PCR products generated with primers SmFwd + CEKG4 (top row) or Sp347 + CEKG4 (bottom row), as described in the Methods. In both rows, lanes 1-10 represent ST-PCR reactions from 10 colonies of MDS30 after IS*3*::SpR transposition. Lanes 11 and 12 represent ST-PCR reactions from two colonies of MDS42 after IS*3*::SpR transposition. Lanes 13 and 14 are negative control ST-PCRs, made on MDS30 and MDS42 colonies, and lane 15 is a positive control ST-PCR made on MDS30_IS*3*::SpR. Yellow rectangles mark the ST-PCR products chosen for sequencing. M: GeneRuler 1 kb Plus DNA Ladder (Thermo Scientific)
