## Supplement tables for "inPOSE: a flexible toolbox for chromosomal cloning and amplification of bacterial transgenes"

Table S1. Primers used in this study

| **name** | **sequence** |
| --- | --- |
| Km13fw | GTCAAGAAGGCGATAGAAGGCG |
| Km96rev | GGCGAATGGGCTGACCG |
| Sp347fw | GCTGGACCTACCAAGGCAAC |
| Sp439rev | TGCAGGTATCTTCGAGCCAG |
| lacZ110fw | CCGTCGATATTCAGCCATGTGC |
| lacZ200rev | TCAGCCGCTACAGTCAACAG |
| Sp110fw | CTAGCTTCAAGTATGACGGGCTG |
| IS1D4 | aagccttacgcgaagcga |
| YeaE3 | ACGCTCCTGAGATTACA |
| IS1ASpF | GATGGTGTTTTTGAGGTGCTCCAGTGGCTTCTGTTTC*TATTTAACGACCCTGCCCTG* |
| IS1ASpR | TGACTTTGTCATGCAGCTCCACCGATTTTGAGAACGACAGC*AAGGGAGAAAGGCGGAACAG* |
| IS1ACmR_F | ATGGTGTTTTTGAGGTGCTCCAGTGGCTTCTGTTTCTAT*AACAGGTTGAACTGCGGATC* |
| IS1ACmR_R | TGACTTTGTCATGCAGCTCCACCGATTTTGAGAACGACAG*CCTGTGACGGAAGATCACTTCG* |
| IS1Sp100F | CCAGTGGCTTCTGTTTCTATCAGCTGTCCCTCCTGTTCAGCTACTGACGGGGTGGTGCGTAACGGCAAAAGCACCGCCGGACATCAGCGCTATCTCTGCTTATTTAACGACCCTGCCCTG |
| IS1Sp100R | ACTTTGTCATGCAGCTCCACCGATTTTGAGAACGACAGCGACTTCCGTCCCAGCCGTGCCAGGTGCTGCCTCAGATTCAGGTTATGCCGCTCAATTCGCAACTTTGTATGTGTCCGCAGC |
| IS3SpF | TCAGAGGTGACTCACATGACAAAAACAGTATCAACCAGTACTATTTAACGACCCTGCCCTG |
| IS3SpR | CACACCAACTGTGCCGCCGCCACCGATTGTAATCACATTCGACCGAGTGAGCTGGCTATTTG |
| IS3flanking1 | CCTAAATTAGCGCCCGTTCC |
| IS3flanking2 | TTAATCGTGGTGTGGTAGAAGG |
| pKdSeq5 | CAGTGAATGGGGGTAAATGGC |
| pKDseq3 | ACCACCGCGCTACTGC |
| pKD1 | tttataacctccttagagctcga |
| IS1noScarFwd | GTGCGTACCGGGTTGAGAAGgttttagagctagaaatagcaag |
| pKD2 | ccaattgtccatattgcatca |
| IS1noScarRev | CTTCTCAACCCGGTACGCACgtgctcagtatctctatcactga |
| IS1noScarCheckR | ACCTTCTCAACCCGGTACG |
| IS3noScarFwd | CTGCGACAGCGTTGTCCTCGgttttagagctagaaatagcaag |
| IS3noScarRev | CGAGGACAACGCTGTCGCAGgtgctcagtatctctatcactga |
| IS3noScarCheckR | AACCGAGGACAACGCTG |
| IS1_Ssp_up | aatATTTAGTGTATGATGGTGTTTTTG |
| IS1_Xho_Rev | TCAAACTCGAGTCGCCATAGTGCG |
| IS1_Xho_Fwd | GGCGACTCGAGTTTGACGTGGTGATATGG |
| IS1_Ssp_Dn | aatATTGATAGTGTTTTATGTTCAG |
| pSTA-F | ccagttcgatgtaacccact |
| pSTA-R | ggatctgaggttcttatggct |
| T7 universal | taatacgactcactataggg |
| IS3_Ssp_up | aatATTCGCCTGAATTTCGC |
| IS3_Xho_Rev | cCTCGAGCGCAGATTATGCCGC |
| IS3_Xho_Fwd | tctgcgCTCGAGGTTGCTGCTACG |
| IS3_Ssp_Dn | aatATTGTTCCGGACTGAG |
| IS1VioFwd | CAATCTGCTCTGATGCCGCACGAGTTTGACGTGGTGATATGG |
| IS1VioRev | CAGCTCATTTCTTAAGTCTCCCGACTGCGGCCTGAG |
| pUTLIQfwd | GAGACTTAAGAAATGAGCTG |
| VioCendRev2 | GCGCCTGTTTCAGTTGGCTGG |
| VioCendfw | GGTACAAGATAGGGAGG GTCAAC |
| pUTLIQRev7 | AGTCTTTCGACTGAGCCTTTC |
| VioARev | GTGAACGGATACACCTC GCTC |
| IS3VioFwd | CAATCTGCTCTGATGCCGCACGAGGTTGCTGCTACGATAATG |
| IS3VioRev | CAGCTCATTTCTTAAGTCTCATCCCGTTCTGCCAGC |
| SmFw | CTTACGTTGTCCCGCATTTGG |
| SmRev | ATATCACTGTGTGGCTTCAGG |
| TetR1 | cactagagaacatactggcta |
| pSTKF | GTCTGTTGTGCCCAGTCATAG |
| Km_HindIII_Fwd | GGAAGCTT-AGGATCGTTTCGCATGATTG |
| Km_HindIII_Rev | GGAAGCTT-GACGGTATCGAACCCCAGAG |
| pCas9Fwd | CAGCTAGGAGGTGACTGAAG |
| pCas9Rev | GGACGATCACACTACTCTTC |
| IS1_SPC_Chek_Rev | CTTCTCAACCCGGTACGCAC |
| IS1-SPC1 | AAACTGATTTTCTGGTGCGTACCGGGTTGAGAAGG |
| IS1-SPC2 | AAAACCTTCTCAACCCGGTACGCACCAGAAAATCA |
| SPC-IS3-Fwd-long | aaacTCCGCCAACACTGCGACAGCGTTGTCCTCGg |
| SPC-IS3-Rev-long | aaaacCGAGGACAACGCTGTCGCAGTGTTGGCGGA |
| SPC-IS3-Fwd-short | aaacCTGCGACAGCGTTGTCCTCGg |
| SPC-IS3-Rev-short | aaaacCGAGGACAACGCTGTCGCAG |
| IS3_SPC_Chek_Rev | gctctaaaacCGAGGACAACG |
| InsAfw | cgcATGctcGTGGCTTCTGTTTCTATCAG |
| InsArev | gcttcttcAAGTGACGTAAAATCGTG |
| InsBfw | CGTCACTTgaagaagCTCAGGCCGCAG |
| InsBrev | ttcggatccTTATTGATAGTGTTTTATGTTCAG |
| pZA31rev | TACGCATGCGGTACCTT |
| pZA31fw | ctaggatccATAAGCTTAATTAGCTGAGTCTAGAG |
| tetP5 | ACTGAGCACATCAGCAGGAC |
| IS1/1 | tgagaacgacagcgac |
| pSToriF | acctagGCTTGGCACTGGCTGATC |
| pSToriR | gattaatACAGCGTTTGCGACATCC |
| tnp3rev | ACGCAGAACACGGCACATAG |
| pZArev | TCTAGGGCGGCGGATTTGTC |
| Ecob1028fw | tctgaattcaccgggcgtaataaggtgg |
| Bamb1028rev | tgtggaTCCAGTCCGGCAAGATAATCG |
| pSGR | CACATACGATTTAGGTGACACT |
| Ap1 | cctccatccagtctattaattgtt |
| CEKG2B: | GGCCACGCGTCGACTAGTACNNNNNNNNNNACGCC |
| CEKG4: | GGCCACGCGTCGACTAGTAC |
| SpRVioIntFw | *AGAGAAGATTTTCAGCCTGATACAGATTAAATCAGAACGC*CTATTTAACGACCCTGCCCTG |
| SpRVioIntRev | *TTTCGTTTTATTTGATGCCTGGCAGTTCCCTACTCTCGCA*CCGAGTGAGCTGGCTATTTG |
| KmRVioIntFw | *AGAGAAGATTTTCAGCCTGATACAGATTAAATCAGAACGC*AGCCATGAGGGTTTAGTTCG |
| KmRVioIntRev | *TTTCGTTTTATTTGATGCCTGGCAGTTCCCTACTCTCGCA*CCTGTTATCCCTAGCGGATC |

Table S2. The efficiencies of recombineering with linear DNA fragments

| Strain | IS element tpe | no of IS elements in genome | linear DNA type | ng DNA used | total colonies per ml | PCR positive colonies | % of positive colonies | calculated total correct colonies | absolute efficiency of integration (recombinant/ ng DNA) |
| --- | --- | --- | --- | --- | --- | --- | --- | --- | --- |
| MDS42.IS1 | IS1 | 1 | IS1::SpR/short homology | 100 | 2064 | 10 out of 10 | 100 | 2064 | 20.64 |
| MDS42.IS1 | IS1 | 1 | IS1::SpR/long homology | 100 | 26880 | 10 out of 10 | 100 | 26880 | 268.80 |
| MDS39R | IS1 | 2 | IS1::SpR/short homology | 100 | 31300 | 5 out of 10 | 50 | 15650 | 156.50 |
| MDS30 | IS3 | 1 | IS3::SpR | 100 | 920 | 10 out of 10 | 100 | 920 | 9.20 |
| BLK09 | IS3 | 2 | IS3::SpR | 100 | 272 | 10 out of 10 | 100 | 272 | 2.72 |
| MDS27 | IS3 | 2 | IS3::SpR | 100 | 860 | 10 out of 10 | 100 | 860 | 8.60 |
| MDS42.IS1 | IS1 | 1 | IS1::VioAESpR | 450 | 2400 | 6 out of 8 | 75 | 1800 | 4.00 |
| MDS42.IS1 | IS1 | 1 | IS1::VioAEKmR | 400 | 1010 | 20 out of 20 | 100 | 1010 | 2.53 |
| MDS39R | IS1 | 2 | IS1::VioAESpR | 450 | 1690 | 7 out of 20 | 35 | 591.5 | 1.31 |
| MDS39R | IS1 | 2 | IS1::VioAEKmR | 400 | 3400 | 13 out of 20 | 65 | 2210 | 5.53 |
| MDS30 | IS3 | 1 | IS3::VioAESpR | 1000 | 10 | 3 out of 10 | 30 | 3 | 0.00 |
| MDS30 | IS3 | 1 | IS3::VioAEKmR | 750 | 4060 | 10 out of 10 | 100 | 4060 | 5.41 |
| BLK09 | IS3 | 2 | IS3::VioAESpR | 1000 | 5090 | 4 out of 20 | 20 | 1018 | 1.02 |
| BLK09 | IS3 | 2 | IS3::VioAEKmR | 750 | 630 | 13 out of 20 | 65 | 409.5 | 0.55 |
| BLK16 | IS3 | 2 | IS3::VioAESpR | 1000 | 3820 | 15 out of 20 | 75 | 2865 | 2.87 |
| BLK16 | IS3 | 2 | IS3::VioAEKmR | 750 | 3350 | 10 out of 20 | 50 | 1675 | 2.23 |
| MDS27 | IS3 | 2 | IS3::VioAESpR | 1000 | 450 | 5 out of 20 | 25 | 112.5 | 0.11 |
| MDS27 | IS3 | 2 | IS3::VioAEKmR | 750 | 900 | 9 out of 20 | 45 | 405 | 0.54 |
| MDS42.IS1 | IS1 | 1 | IS1:SpR | 107.6 | 1642 | 10 out of 10 | 100 | 1642 | 15.26 |
| MDS42.IS1 | IS1 | 1 | IS1:CmR | 123.8 | 89 | 10 out of 10 | 100 | 89 | 0.72 |
| MDS39R | IS1 | 2 | IS1:SpR | 107.6 | 2160 | 5 out of 6 | 83.33 | 1799.93 | 16.73 |
| MDS39R | IS1 | 2 | IS1:CmR | 123.8 | 81 | 3 out of 6 | 49.99 | 40.49 | 0.33 |

Table S3. The efficiencies of CRISPR/Cas-mediated recombineering with linear DNA fragments

| Strain | IS element tpe | no of IS elements in genome | System | linear DNA type | ng DNA used | total colonies per ml | PCR-verified single positive colonies | PCR-verified double positive colonies | % of colonies positive by PCR | calculated total correct colonies | Absolute efficiency of integration (recombinant /ng DNA) |
| --- | --- | --- | --- | --- | --- | --- | --- | --- | --- | --- | --- |
| MDS42.IS1 | IS1 | 1 | NO-SCAR-IS1 | IS1:KmR | 100 | 87 | 10 out of 10 | NA | 100.00 | 87.00 | 0.87 |
| MDS42.IS1 | IS1 | 1 | NO-SCAR-IS1 | IS1:KmR | 100 | 97 | 10 out of 10 | NA | 100.00 | 97.00 | 0.97 |
| MDS30 | IS3 | 1 | NO-SCAR-IS3 | IS3:KmR | 200 | 286 | 10 out of 10 | NA | 100.00 | 286.00 | 1.43 |
| MDS30 | IS3 | 1 | NO-SCAR-IS3 | IS3:KmR | 200 | 1114 | 4 out of 10 | NA | 40.00 | 445.60 | 2.23 |
| MDS42.IS1 | IS1 | 1 | pORTMAGE-2/pCas9-IS1 | IS1:SpR | 100 | 5360 | 10 out of 10 | NA | 100.00 | 5360.00 | 53.60 |
| MDS42.IS1 | IS1 | 1 | pORTMAGE-2/pCas9-IS1 | IS1:SpR | 100 | 9160 | 10 out of 10 | NA | 100.00 | 9160.00 | 91.60 |
| MDS30 | IS3 | 1 | pORTMAGE-2/pCas9-IS3 | IS3:SpR | 100 | 800 | 10 out of 10 | NA | 100.00 | 800.00 | 8.00 |
| MDS30 | IS3 | 1 | pORTMAGE-2/pCas9-IS3 | IS3:SpR | 100 | 730 | 10 out of 10 | NA | 100.00 | 730.00 | 7.30 |
| MDS42.IS1 | IS1 | 1 | NO-SCAR-IS1 | IS1:GFP | 100 | 636000 | 3 out of 30 | NA | 10.00 | 63600.00 | 636.00 |
| MDS42.IS1 | IS1 | 1 | NO-SCAR-IS1 | IS1:GFP | 100 | 426000 | 2 out of 65 | NA | 3.08 | 13107.69 | 131.08 |
| MDS42.IS1 | IS1 | 1 | NO-SCAR-IS1 | IS1:GFP | 100 | 410000 | 3 out 0f 50 | NA | 6.00 | 24600.00 | 246.00 |
| MDS30 | IS3 | 1 | pORTMAGE-2/pCas9-IS3 | IS3:GFP | 100 | 5440 | 1 out of 20 | NA | 5.00 | 272.00 | 2.72 |
| MDS30 | IS3 | 1 | pORTMAGE-2/pCas9-IS3 | IS3:GFP | 100 | 6820 | 2 out of 10 | NA | 20.00 | 1364.00 | 13.64 |
| MDS30 | IS3 | 1 | pORTMAGE-2/pCas9-IS3 | IS3:GFP | 100 | 2760 | 3 out of 40 | NA | 7.50 | 207.00 | 2.07 |
| MDS30 | IS3 | 1 | NO-SCAR-IS3 | IS3:VioABCDE-KmR | 600 | 62 | 6 out of 10 | NA | 60.00 | 37.20 | 0.06 |
| MDS42.IS1 | IS3 | 1 | NO-SCAR-IS3 | IS3:VioABCDE-KmR | 750 | 2520 | 15 out of 20 | NA | 75.00 | 1890.00 | 2.52 |
| MDS39R | IS1 | 2 | pORTMAGE-2/pCas9-IS1 | IS1:SpR | 100 | 1780 | NA | 2 out of 10 | 20.00 | 356.00 | 3.56 |
| MDS39R | IS1 | 2 | pORTMAGE-2/pCas9-IS1 | IS1:SpR | 100 | 340 | NA | 3 out of 10 | 30.00 | 102.00 | 1.02 |
| MDS27 | IS3 | 2 | pORTMAGE-2/pCas9-IS3 | IS3:SpR | 100 | 210 | NA | 6 out of 10 | 60.00 | 126.00 | 1.26 |
| BLK16 | IS3 | 2 | pORTMAGE-2/pCas9-IS3 | IS3:SpR | 100 | 1330 | NA | 15 out of 20 | 75.00 | 997.50 | 9.98 |
| MDS27 | IS3 | 2 | NO-SCAR-IS3 | IS3:VioABCDE-KmR | 750 | 400 | NA | 1 out of 13 | 7.60 | 30.40 | 0.04 |
| MDS39R | IS3 | 2 | NO-SCAR-IS3 | IS3:VioABCDE-KmR | 750 | 764 | NA | 1 out of 20 | 5.00 | 38.20 | 0.05 |
